# Osteocytes remodel bone by TGF-β-induced YAP/TAZ signaling

**DOI:** 10.1101/611913

**Authors:** Christopher D. Kegelman, Jennifer C. Coulombe, Kelsey M. Jordan, Daniel J. Horan, Ling Qin, Alexander G. Robling, Virginia. L Ferguson, Teresita M. Bellido, Joel D. Boerckel

**Author notes:** To whom correspondence should be addressed: Joel D. Boerckel.

## Abstract

Osteocytes are bone matrix-entombed cells that form an interconnected network of processes called the lacunar/canalicular system, which enables osteocytes to coordinate bone formation and resorption. Osteocytes indirectly regulate osteoblast and osteoclast activity on bone surfaces but also directly resorb and deposit their surrounding bone matrix through perilacunar/canalicular remodeling. However, the molecular mechanisms by which osteocytes control bone remodeling remain unclear. We previously reported that the transcriptional regulators Yes-associated protein (YAP) and Transcriptional co-activator with PDZ-motif (TAZ) promote bone acquisition in osteoblast-lineage cells. Here, we tested the hypothesis that YAP and TAZ regulate osteocyte-mediated bone remodeling by conditional ablation of both YAP and TAZ from mouse osteocytes using 8kb-DMP1-Cre. Osteocyte conditional YAP/TAZ deletion reduced bone mass and dysregulated matrix collagen content and organization, which together impaired bone mechanical properties. YAP/TAZ deletion reduced osteoblast number and activity and increased osteoclast activity. In addition, YAP/TAZ deletion directly impaired osteocyte lacunar/canalicular network remodeling, reducing canalicular density, length, and branching, but did not alter lacunar size or shape. Further, consistent with recent studies identifying TGF-β signaling as a key inducer of perilacunar/canalicular remodeling through expression of matrix-remodeling enzymes, YAP/TAZ deletion *in vivo* decreased osteocyte expression of matrix proteases Mmp13, Mmp14, and Cathepsin K. *In vitro*, pharmacologic inhibition of YAP/TAZ transcriptional activity in osteocyte-like cells abrogated TGF-β-induced protease gene expression. Together, these data show that YAP and TAZ act downstream of TGF-β in osteocytes to control bone matrix accrual, organization, and mechanical properties indirectly by coordinating osteoblast/osteoclast activity and directly by regulating perilacunar/canalicular remodeling.

## INTRODUCTION

Skeletal fragility diseases are characterized by decreased bone strength. Bone strength is determined by both the quantity and quality of the bone, both of which are necessary to explain fracture susceptibility (1–3). Decreased bone mass is a hallmark of osteoporosis, but defects in bone geometry, microarchitecture, porosity, or matrix material properties are significant contributors to bone fragility (4). Both bone quantity and quality are influenced by bone remodeling, a tightly-coordinated process by which old, damaged bone is resorbed and new, strong bone is deposited. Imbalanced bone remodeling, either by excessive bone resorption or decreased bone formation can impair bone quantity and quality (5, 6). Understanding the mechanisms that control bone remodeling is both scientifically and therapeutically critical.

As the most abundant cell type in bone, osteocytes control bone remodeling both indirectly and directly (7). As terminally differentiated osteoblasts, osteocytes reside within lacunae in the mineralized bone matrix, and extend dendritic processes through an interconnected network of micro-channels called canaliculi that permeate the bone matrix (8–10). In humans, this lacunar/canalicular network contains an estimated 3.7 trillion dendritic projections and covers more than 215 m^2^ of bone surface area (11). Through this widespread network, osteocyte-derived molecules reach the bone surfaces and regulate bone remodeling indirectly by coordinating osteoclast-osteoblast coupled remodeling (12, 13). In addition, osteocytes directly resorb and deposit bone in their surrounding bone matrix via perilacunar/canalicular remodeling (14–16). However, the molecular mechanisms by which osteocytes control bone remodeling remain poorly understood.

Indirect bone remodeling is regulated in part through the CCN family of matricellular growth factors, cysteine-rich angiogenic inducer-61 (Cyr61) and connective tissue growth factor (Ctgf) (17, 18). Cyr61 and Ctgf have been implicated in activation of osteoblastogenesis (19–22) and inhibition of osteoclastogenesis (23). Both Cyr61 and Ctgf are downstream gene targets of the paralogous transcriptional regulators Yes associated protein (YAP) and Transcriptional co-activator with PDZ-binding motif (TAZ) (24, 25). YAP and TAZ are the key effector proteins of the Hippo signaling pathway and regulate important biological functions including organ size determination, tissue regeneration, and cancer (26, 27). Transcriptional complex formation of nuclear YAP and/or TAZ with the transcriptional enhancer activator domain (TEAD) family proteins is required to induce expression of Cyr61 and Ctgf (24, 28). In addition to paracrine regulation of osteoblast/osteoclast activity, osteocytes also directly remodel the bone matrix through perilacunar/canalicular remodeling, which is regulated by TGF-β(29). The TGF-β and YAP/TAZ signaling pathways are known to interact in a variety of cell types including cancer cells, fibroblasts, and epithelial cells (30–33). These observations position YAP and TAZ as potential mediators of both direct and indirect osteocyte-mediated bone remodeling.

Here, we conditionally ablated both YAP and TAZ from DMP1-expressing cells and evaluated bone remodeling. We found that osteocyte YAP and TAZ promote bone formation *in vivo* by coordinating osteoclast/osteoblast activity and also regulate osteocytic perilacunar/canalicular remodeling. *In vitro*, YAP/TAZ inhibition abrogated TGF-β-induced expression of growth factors and matrix proteases that mediate both osteocyte-indirect and -direct bone remodeling, respectively. Together, these data position YAP and TAZ downstream of TGF-β signaling in osteocytes to control bone matrix accrual, organization, and mechanical properties by regulating both direct perilacunar/canalicular remodeling and indirect osteoblast/osteoclast activity.

## RESULTS

### DMP1-Cre conditionally ablates YAP and TAZ primarily in osteocytes

To determine the roles of YAP and TAZ in osteocyte-mediated bone remodeling, we used 8kb-DMP1-Cre mice to selectively delete YAP and TAZ from osteocytes (34, 35). We used a breeding strategy that generated YAP/TAZ allele dosage-dependent DMP1-conditional knockouts (36). All genotypes (Table 1) appeared at expected Mendelian ratios. By early skeletal maturity (post-natal day 84, P84), YAP/TAZ allele dosage-dependent DMP1-conditional deletion did not significantly alter body mass in either males or females (Fig. S1A). YAP/TAZ deletion reduced femoral length at P84 only in double homozygous knockouts, for both sexes (Fig. S1B, Fig. 1A-B). A single copy of either gene was sufficient to rescue this defect, suggesting some degree of functional redundancy. Therefore, for further analyses, we selected littermate YAP^fl/fl^;TAZ^fl/fl^ wild type (YAP^WT^;TAZ^WT^) and YAP^fl/fl^;TAZ^fl/fl^;8kb-DMP1-Cre conditional double knockout (YAP^cKO^;TAZ^cKO^) mice for comparison.

**Table 1:** Experimental genotypes & abbreviations.

| Genotype | Abbreviation |
| --- | --- |
| Yap<sup>fl/fl</sup>;Taz<sup>fl/fl</sup> | YAP<sup>WT</sup>;TAZ<sup>WT</sup> |
| Yap<sup>fl/+</sup>;Taz<sup>fl/+</sup>;DMP1<sup>cre/+</sup> | YAP<sup>cHET</sup>;TAZ<sup>cHET</sup> |
| Yap<sup>fl/fl</sup>;Taz<sup>fl/+</sup>;DMP1<sup>cre/+</sup> | YAP<sup>cKO</sup>;TAZ<sup>cHET</sup> |
| Yap<sup>fl/+</sup>;Taz<sup>fl/fl</sup>;DMP1<sup>cre/+</sup> | YAP<sup>cHET</sup>;TAZ<sup>cKO</sup> |
| Yap<sup>fl/fl</sup>;Taz<sup>fl/fl</sup>;DMP1<sup>cre/+</sup> | YAP<sup>cKO</sup>;TAZ<sup>cKO</sup> |

**Figure 1.**
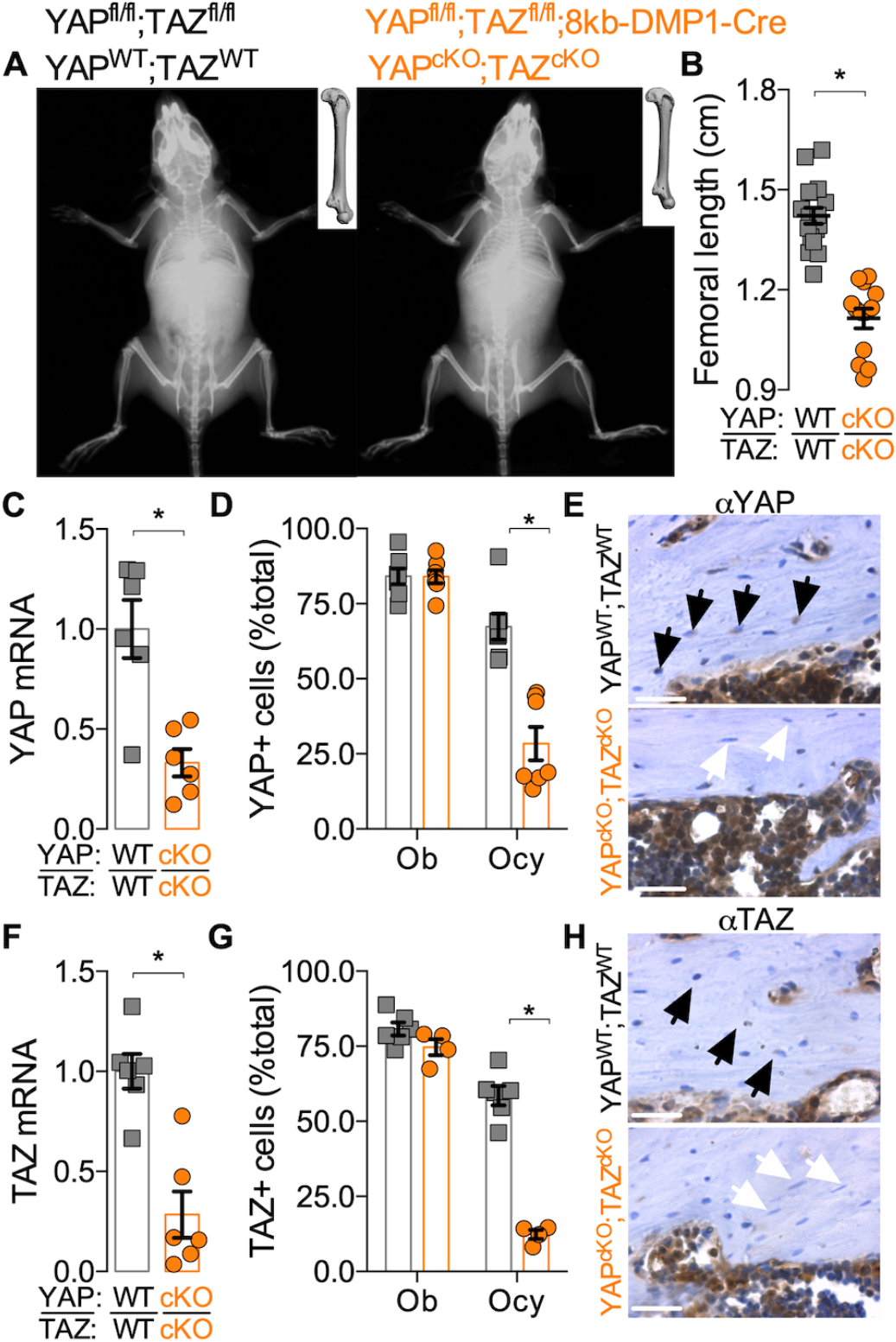
8kb-DMP1-Cre selectively ablated YAP/TAZ expression from osteocytes. **A)** Representative radiographs for wild type (YAP^WT^;TAZ^WT^) and conditional double knockout (YAP^cKO^;TAZ^cKO^) mice at P84. Insets: femur microCT reconstructions. **B)** Femoral lengths measured at P84. **C-H)** Recombination efficiency and specificity was assessed by measurement of YAP and TAZ mRNA and protein expression. **C)** YAP mRNA expression, relative to 18s rRNA, from femoral bone preparations at P84. **(D)** YAP protein expression from femoral sections at P28. **E)** Representative micrographs of YAP immunostaining in YAP^WT^;TAZ^WT^ and YAP^cKO^;TAZ^cKO^ femurs at P28. **F)** TAZ mRNA expression, relative to 18s rRNA, from femoral bone preparations at P84. **(G)** TAZ protein expression from femoral sections at P28. **H)** Representative micrographs of TAZ immunostaining in WT and YAP^cKO^;TAZ^cKO^ femurs at P28. N = 8 per group for qPCR and N = 4-6 per group for IHC. Black arrows indicate positively-labeled osteocytes. White arrows indicate negative osteocytes. Scale bars equal 50 μm.

Recent reports suggest that both the 8 kb and 10 kb DMP1 promoter fragments used to drive Cre recombinase expression for conditional ablation in osteocytes can also induce recombination in mature osteoblasts, depending on the sensitivity of the floxed alleles (35, 37–39). Therefore, to assess the specificity of 8kb-DMP1-Cre-mediated YAP/TAZ deletion, we evaluated YAP/TAZ expression in both osteocytes and osteoblasts. YAP mRNA expression was significantly reduced by 67% in YAP^cKO^;TAZ^cKO^ femoral bone preparations (Fig. 1C). YAP protein expression was significantly reduced in osteocytes (58% reduction), but not osteoblasts (0% reduction; p = 0.98) (Fig. 1D-E). Similarly, TAZ mRNA expression was significantly reduced by 72% in YAP^cKO^;TAZ^cKO^ femoral bone preparations (Fig. 1F), and TAZ protein expression was significantly reduced in osteocytes (79% reduction), but not osteoblasts (6% reduction; p = 0.12) (Fig. 1G-H).

### DMP1-conditional YAP/TAZ deletion impaired bone accrual

Dual homozygous YAP/TAZ deletion from DMP1-expressing cells reduced bone accrual and microarchitectural parameters in both the cancellous and cortical compartments of P84 femurs. In metaphyseal cancellous bone, conditional YAP/TAZ deletion significantly reduced bone volume fraction, trabecular number, and thickness and increased trabecular spacing and structural model index (Fig. 2A-F). In mid-diaphyseal femoral cortical bone, conditional YAP/TAZ deletion reduced cortical thickness and bone area without changing medullary area (Fig. 2G-I; Fig. S2). Differences in tissue mineral density (p = 0.24), periosteal perimeter (p = 0.15), and moment of inertia (p = 0.08) were not statistically significant (Fig. 2J-L).

**Figure 2.**
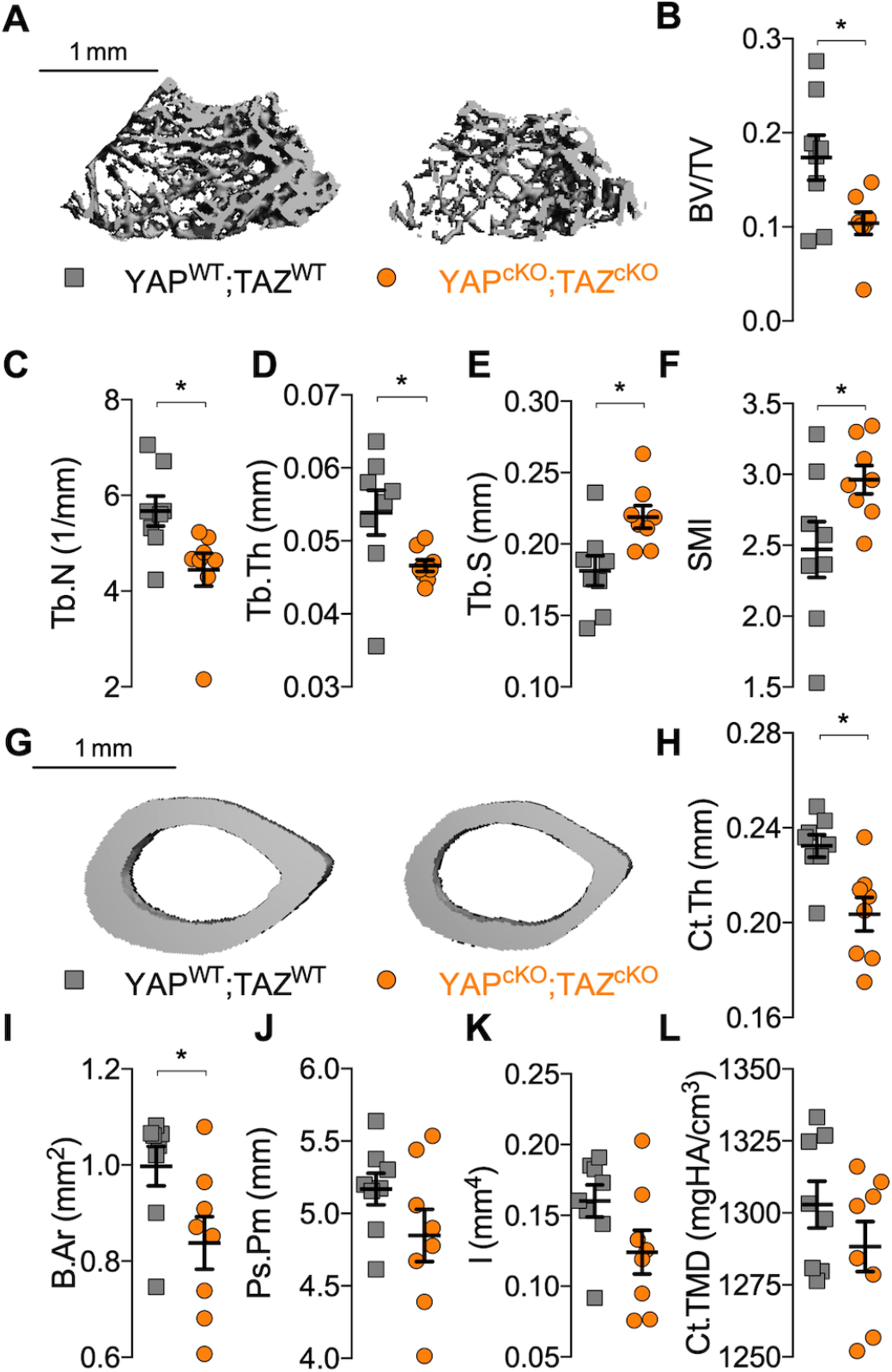
YAP/TAZ ablation from DMP1-expressing cells altered bone microarchitecture. **A)** Representative microCT reconstructions of distal metaphysis of P84 femurs. Quantification of cancellous bone microarchitecture: **(B)** bone volume fraction (BV/TV), **(C)** trabecular thickness (Tb.Th), **(D)** number (Tb.N), **(E)** spacing (Tb.Sp), and **(F)** structural model index (SMI). **G)** Representative microCT reconstructions of mid-diaphysis cortical microarchitecture in P84 femurs. Quantification of cortical microarchitectural properties: **(H)** cortical thickness (Ct.Th), (**I**) bone area (B.Ar), **(J)** periosteal perimeter (Ps.Pm), **(K)** moment of inertia in the direction of bending (I), and **(L)** cortical tissue mineral density (Ct.TMD). Data are presented as individual samples with lines corresponding to the mean and standard error of the mean (SEM). Sample sizes, N = 8. Scale bars indicate 1 mm for microCT reconstructions. These data were originally presented in (40) but removed from the final publication (36).

To determine if the low bone mass phenotype resulted from decreased bone formation and/or increased bone resorption, we evaluated osteoblast and osteoclast activity. Conditional YAP/TAZ deletion increased osteoclast surface per bone surface (Fig. 3A-B) and decreased osteoblast number per bone surface (Fig. 3C-D) in P84 distal femur metaphyseal cancellous bone. YAP/TAZ deletion decreased bone formation rate, reducing both mineralizing surface percentage and mineral apposition rate at P28 in distal femur metaphyseal cancellous bone (Fig. 3E-H). In the cortical compartment, YAP/TAZ deletion similarly reduced osteoblast number per endosteal bone surface (Fig. S3A-B) and decreased bone formation rate by reducing mineral apposition rate without altering mineralizing surface percentage (Fig. S3C-F).

**Figure 3.**
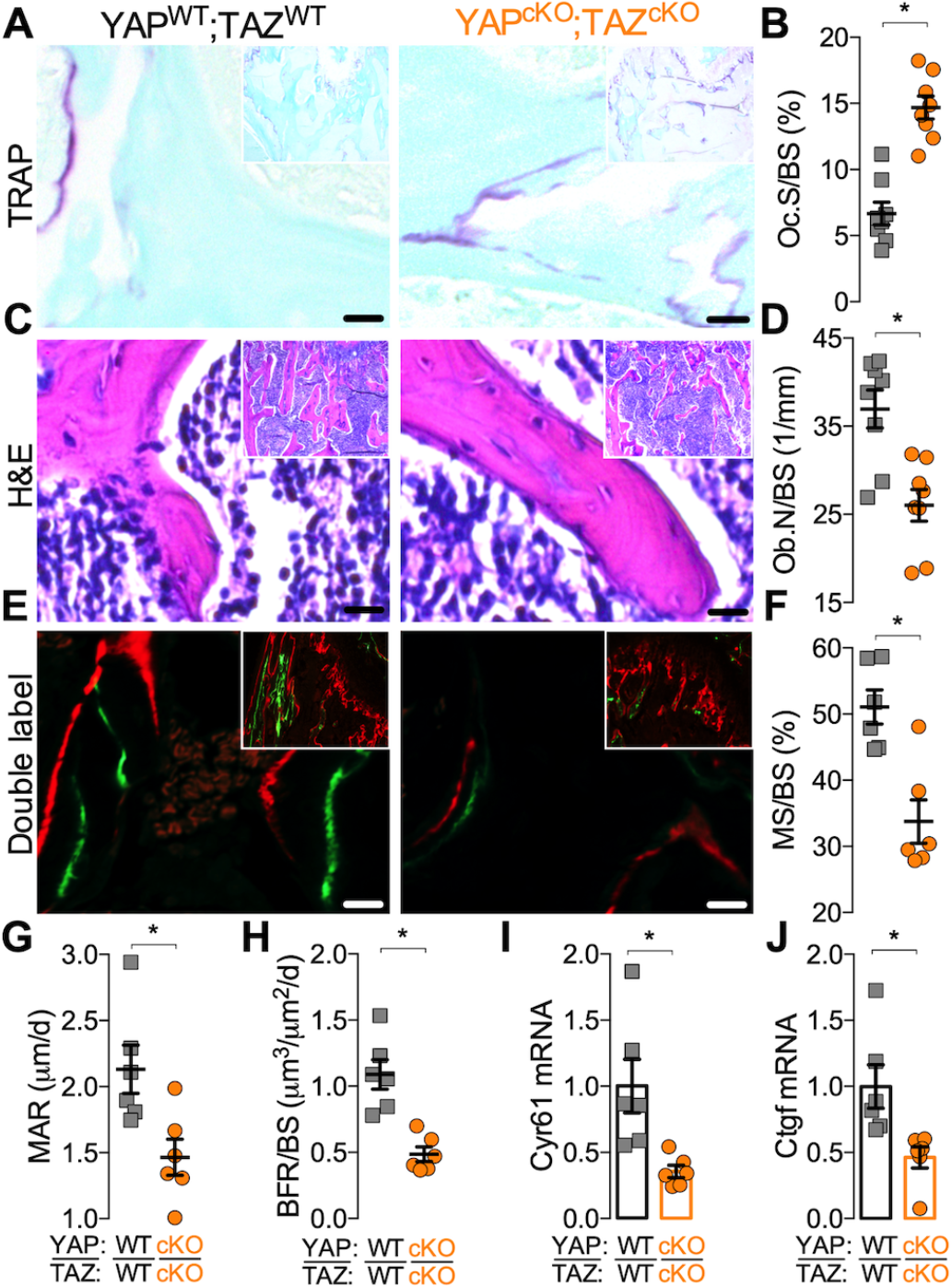
YAP/TAZ ablation from DMP1-expressing cells increased osteoclast activity and decreased osteoblast activity. **A)** Representative high magnification micrographs with insets of low magnification micrographs of P84 cancellous metaphyseal bone stained by TRAP. **B)** Quantification of osteoclast surface per bone surface (Oc.S/BS). **C)** Representative high magnification micrographs with insets of low magnification micrographs of P84 cancellous metaphyseal bone stained by H+E. **D)** Quantification of osteoblast number per bone surface (Ob.N/BS). **E)** Representative high magnification micrographs with insets of low magnification micrographs of double flourochrome labeled P28 cancellous metaphyseal bone. **F-H)** Quantification of **(F)** mineralizing surface percentage (MS/BS), **(G)** mineral apposition rate (MAR) and **(H)** bone formation rate (BFR/BS). **I-J)** Femoral bone preparations from P84 mice were harvested to quantify mRNA expression. Expression levels, normalized to 18S rRNA, were evaluated for **(I)** cysteine-rich angiogenic inducer 61 (Cyr61) and **(J)** connective tissue growth factor (Ctgf). Data presented as individual samples with lines corresponding to the mean and standard error of the mean (SEM). Sample sizes N = 6-8. Scale bars indicate 30 μm in all images.

Further, consistent with our initial hypothesis that YAP/TAZ coordinate osteoblast/osteoclast activation by regulating Cyr61 and Ctgf expression, YAP/TAZ deletion significantly reduced mRNA expression of YAP/TAZ-TEAD target genes, *Cyr61* and *Ctgf* in femoral bone at P84 (Fig. 3I,J). However, YAP/TAZ deletion did not alter sclerostin (*SOST*), receptor activator of nuclear factor kappa-B ligand (*Rankl*) or osteoprotegerin (*Opg*) transcript expression (Fig. S4C-E).

### DMP1-conditional YAP/TAZ deletion impaired bone mechanical properties and matrix collagen composition

To determine whether YAP/TAZ deletion impaired functional mechanical properties, we tested P84 femurs in three point bending to failure (Fig. 4A). Conditional YAP/TAZ deletion did not significantly reduce maximum load to failure (Fig. 4B; p=0.07) but significantly reduced bending stiffness (Fig. 4C; p < 0.05). YAP/TAZ deletion reduced work to maximum load and work to failure (Fig. 4D-E).

**Figure 4.**
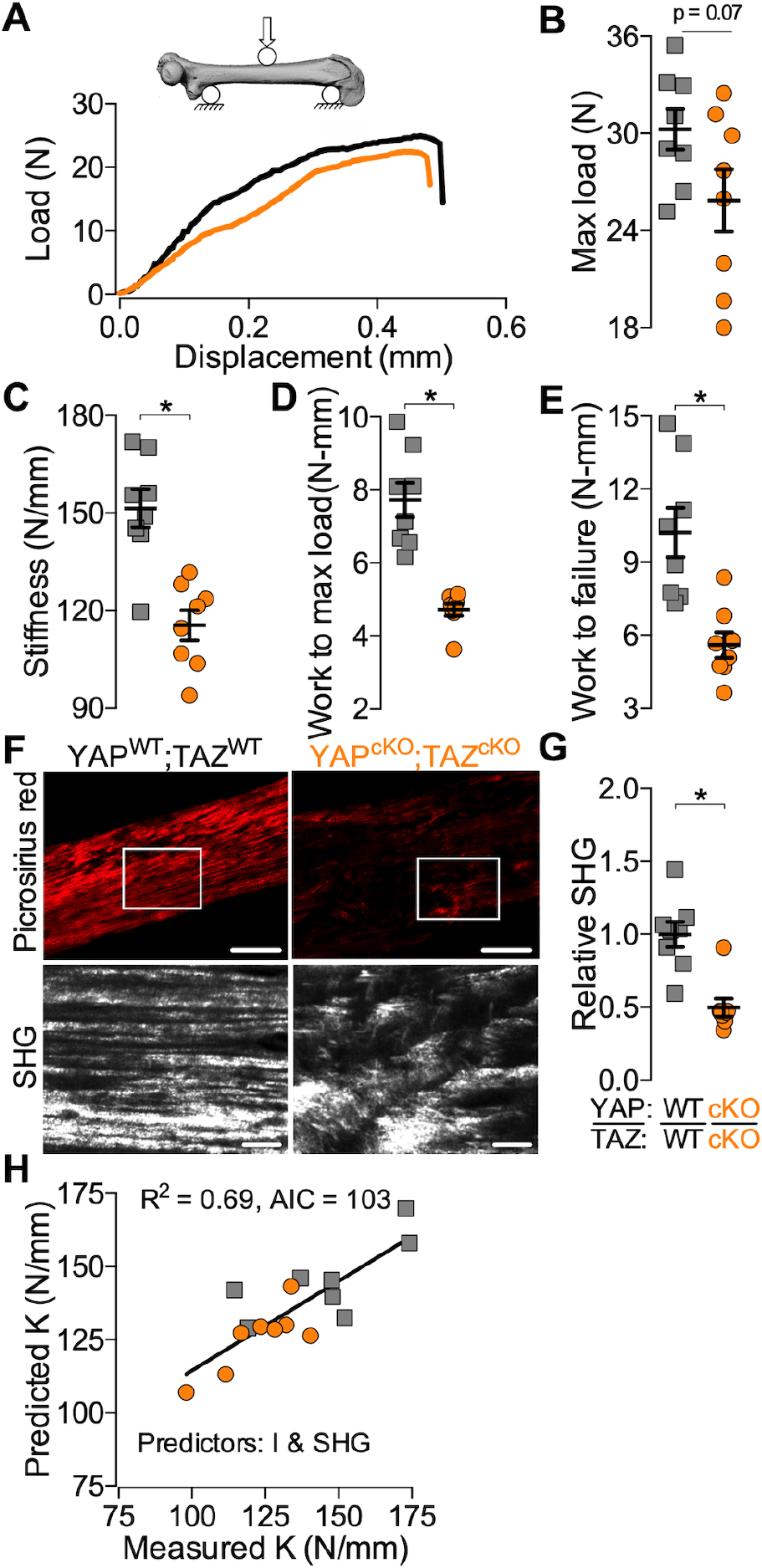
YAP/TAZ ablation from DMP1-expressing cells impaired bone mechanical behavior by altering bone geometry and matrix collagen. **A)** P84 femurs were tested in three-point bending to failure. **B-E)** Quantification of **(B)** maximum load to failure, **(C)** bending stiffness, **(D)** work to max load, and **(E)** work to failure. **F)** Representative micrographs of polarized light microscopy and second harmonic generated (SHG) images of cortical bone from the P84 femurs tested in three-point bending. **G-H)** Quantification of **(G)** SHG intensity normalized to wild type. **H)** Best-subset multivariate regression model predicting experimental bending stiffness (K) using moment of inertia and SHG intensity as predictors. Data presented as individual samples with lines corresponding to the mean and standard error of the mean (SEM). Sample sizes, N = 8 per group. Scale bars equal 100 and 25 μm in the Picrosirius red and SHG, respectively. Portions of these data were originally presented in (40) but removed from the final publication (36).

Intrinsic bone material properties are determined, in part, by collagen content and organization (36). To test whether osteocyte YAP/TAZ regulate the local collagen matrix, we performed polarized light and second harmonic generation (SHG) microscopy (Fig. 4F). YAP/TAZ deletion significantly reduced SHG intensity per bone area, indicative of collagen content and organization (Fig. 4G). To determine whether collagen content and organization contributed to bone mechanical behavior, we performed a Type II multivariate best-subsets regression analysis (36). Both cross sectional bone geometry (section modulus and moment of inertia) and collagen matrix content and organization significantly contributed to bone mechanical behavior, while microCT-measured tissue mineral density did not (Fig. 4H, Fig. S5A-F).

### DMP1-conditional YAP/TAZ deletion impaired the osteocyte canalicular network, but not lacunar morphology

We initially hypothesized that the primary function of YAP/TAZ in osteocytes was to regulate expression of genes that control osteocyte-osteoblast and -osteoclast communication, such as secreted factors Ctgf and Cyr61. However, the reduced bone mechanical properties and disorganized collagen matrix in mice lacking YAP/TAZ from osteocytes phenocopied mice lacking matrix metalloproteinase-13 (Mmp13) (41). Mice lacking Mmp13 had defective collagen organization as a result of impaired perilacunar/canalicular remodeling (41). This led us to ask whether YAP/TAZ deletion from osteocytes caused osteocyte-intrinsic defects, particularly in perilacunar/canalicular remodeling.

Conditional YAP/TAZ deletion did not significantly alter osteocyte density but increased empty lacuna percentage in cancellous metaphyseal bone (Fig. 5A-C). Accordingly, conditional YAP/TAZ deletion increased the number of TUNEL-positive lacunae, suggesting an increase in osteocyte apoptosis within the cancellous bone compartment (Fig. 5D-E). To determine if conditional YAP/TAZ deletion altered the canalicular network, we performed silver nitrate staining and quantified the canalicular processes in 2D (42, 43). Conditional YAP/TAZ deletion significantly reduced canalicular density and mean process length (Fig. 5F-H).

**Figure 5.**
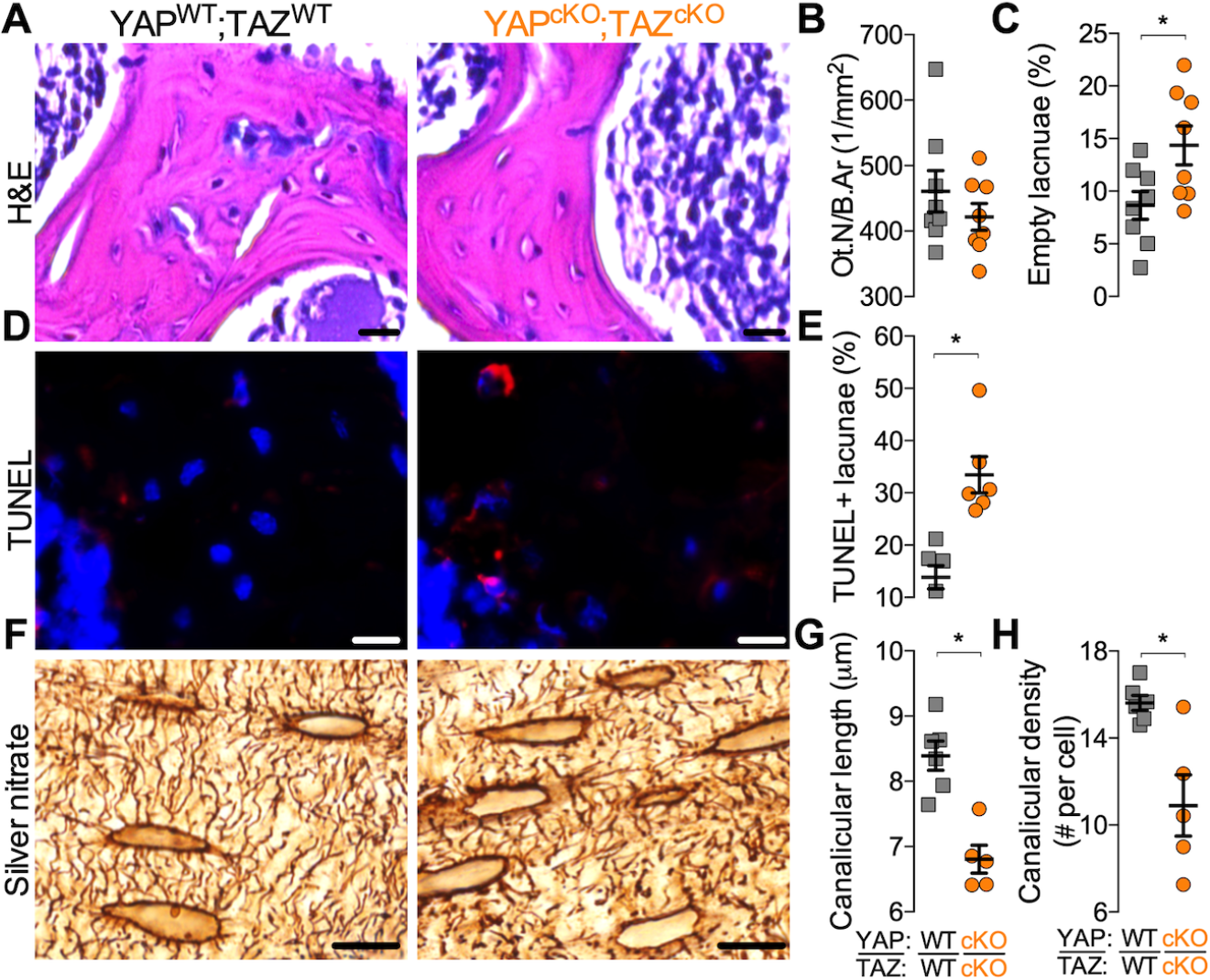
YAP/TAZ ablation from DMP1-expressing cells increased osteocyte apoptosis and reduced canalicular number and length in 2D. **A)** Representative micrographs of P84 cancellous metaphyseal bone stained by H+E. **B-C)** Quantification of **(B)** osteocyte number per bone area (Ot.N/B.Ar), and **(C)** percentage of empty lacunae. **D)** Representative immunofluorescence micrographs of P28 cancellous metaphyseal bone stained with TUNEL positive cells (red) and DAPI (blue). **E)** Quantification of the percentage of TUNEL positive lacunae. **F)** Representative micrographs of P84 cortical bone silver stained for the osteocyte canalicular network. **G-H)** Quantification of **(G)** canalicular density per cell and **(H)** average canalicular length. Data presented as individual samples with lines corresponding to the mean and standard error of the mean (SEM). Sample sizes, N = 6-8 per group. Scale bars equal 30, 25, and 20 μm in the H+E, TUNEL, and silver stain, respectively.

The canalicular system is a three-dimensional network of branched cell processes that physically connect osteocytes to their neighbors (11). To quantify these 3D canalicular networks, we stained osteocyte cytoskeletons in 15-micron-thick cortical bone sections with Alexa Fluor 488-conjugated phalloidin and generated 3D reconstructions by laser scanning confocal microscopy (Fig. 6A). YAP/TAZ deletion significantly reduced branch length (Fig. 6B,C), number of branches per cell (Fig. 6D), and number of junctions per cell (Fig. 6E), with effect sizes consistent with 2D canalicular morphologies measured in silver-stained sections (cf. Fig. 5F-H).

**Figure 6.**
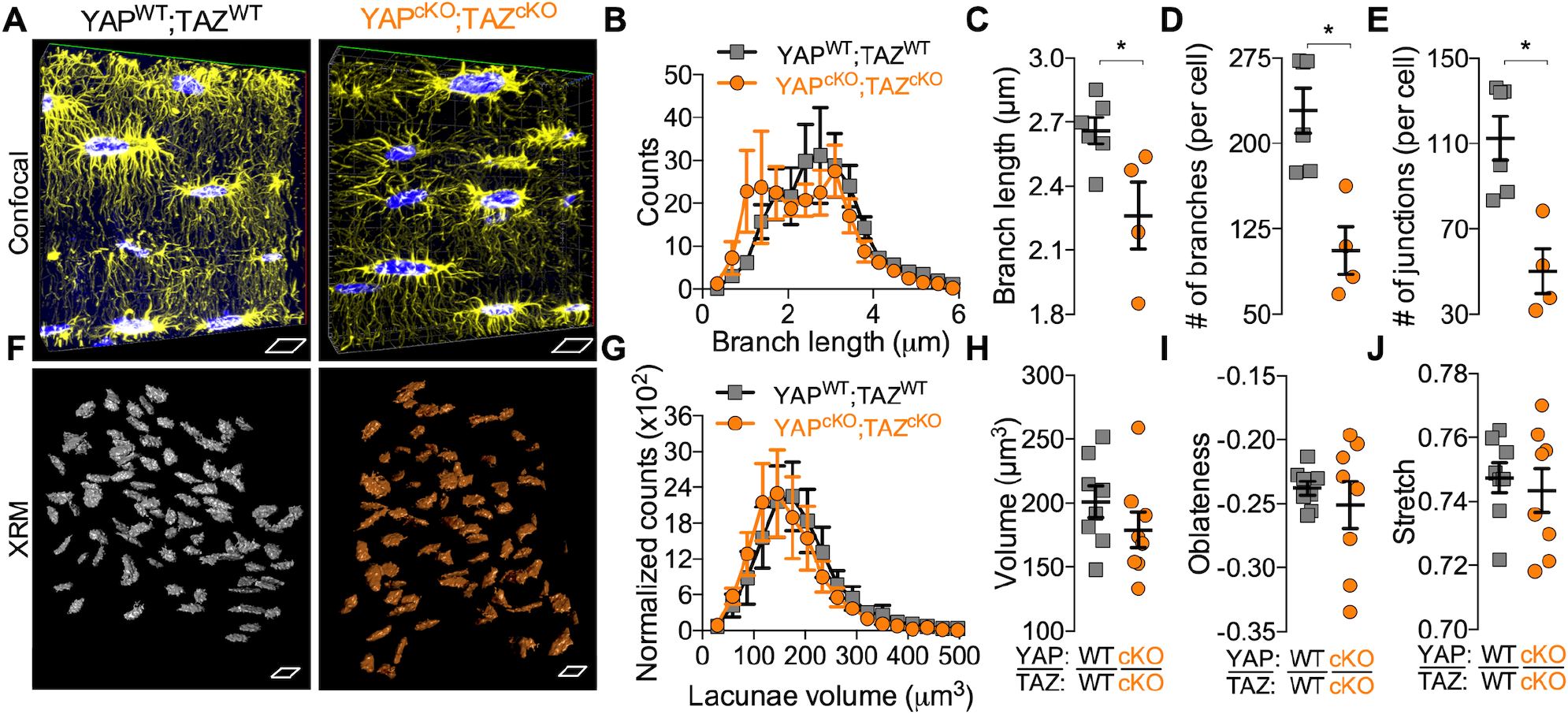
YAP/TAZ ablation from DMP1-expressing cells reduced canalicular network length and branching, but not lacunar morphology in 3D. **A)** Representative 15-micron thick confocal reconstructions of phalloidin-stained osteocyte F-actin cytoskeletons in cortical bone at P28. **B)** Average distributions of branch length. **C-E)** Quantification of **(C)** average branch length, **(D)** number of branches, and **(E)** number of junctions. **F)** Representative lacunar morphologies in 3D in cancellous bone from P84 tibiae using X-ray microscopy (XRM). **G)** Average distributions of lacunar volume. **H-J)** Quantification of **(H)** average lacuna volume, **(I)** oblateness, and **(J)** stretch. Data presented as individual samples with lines corresponding to the mean and standard error of the mean (SEM). Sample sizes, N = 4-6 for confocal network analysis and N = 8 for XRM. Each side of scale parallelograms equal 10 μm for both confocal and XRM.

To determine the effect of YAP/TAZ deletion on osteocyte lacunar morphology, we performed high resolution (0.6 um voxel) X-Ray microscopy (XRM) imaging of both cortical and cancellous bone in the proximal tibia metaphysis (Fig. 6F). Both sex and bone compartment (i.e., cortical vs. cancellous) were statistically significant predictors of lacuna volume and the shape parameters, oblateness (Eq. 1) and stretch (Eq. 2). However, YAP/TAZ deletion did not significantly alter lacuna volume (Fig. 6G,H) or shape (Fig. 6I,J). Statistically significant interactions were observed between sex, bone compartment, and genotype for all lacunar morphology parameters.

### DMP1-conditional YAP/TAZ deletion reduced osteocyte-mediated bone matrix remodeling

Perilacunar/canalicular remodeling involves both the deposition and degradation of the bone matrix directly surrounding the osteocytes. Prior studies used ^3^H-proline pulses and tetracycline labeling to identify osteocyte lacunae as sites of active collagen deposition and mineralization (14, 44, 45).

Here, we measured collagen I gene expression and flourochrome incorporation in osteocyte lacunae to assess peri-osteocyte matrix deposition. Conditional YAP/TAZ deletion reduced the percentage of Calcein-labeled osteocyte lacunae (Fig. 7A,B) and reduced transcript expression of Col1a1 (Fig. 7C). However, YAP/TAZ deletion did not alter expression of either osteogenic genes, including alkaline phosphatase (*Alp*), osteocalcin (*Ocn*), and bone sialoprotein (*Bsp*), or osteocyte-marker genes, dentin matrix protein-1 (*Dmp1*) or phosphate-regulating neutral endopeptidase, X-linked (*Phex*) (Fig. S6A-E).

**Figure 7.**
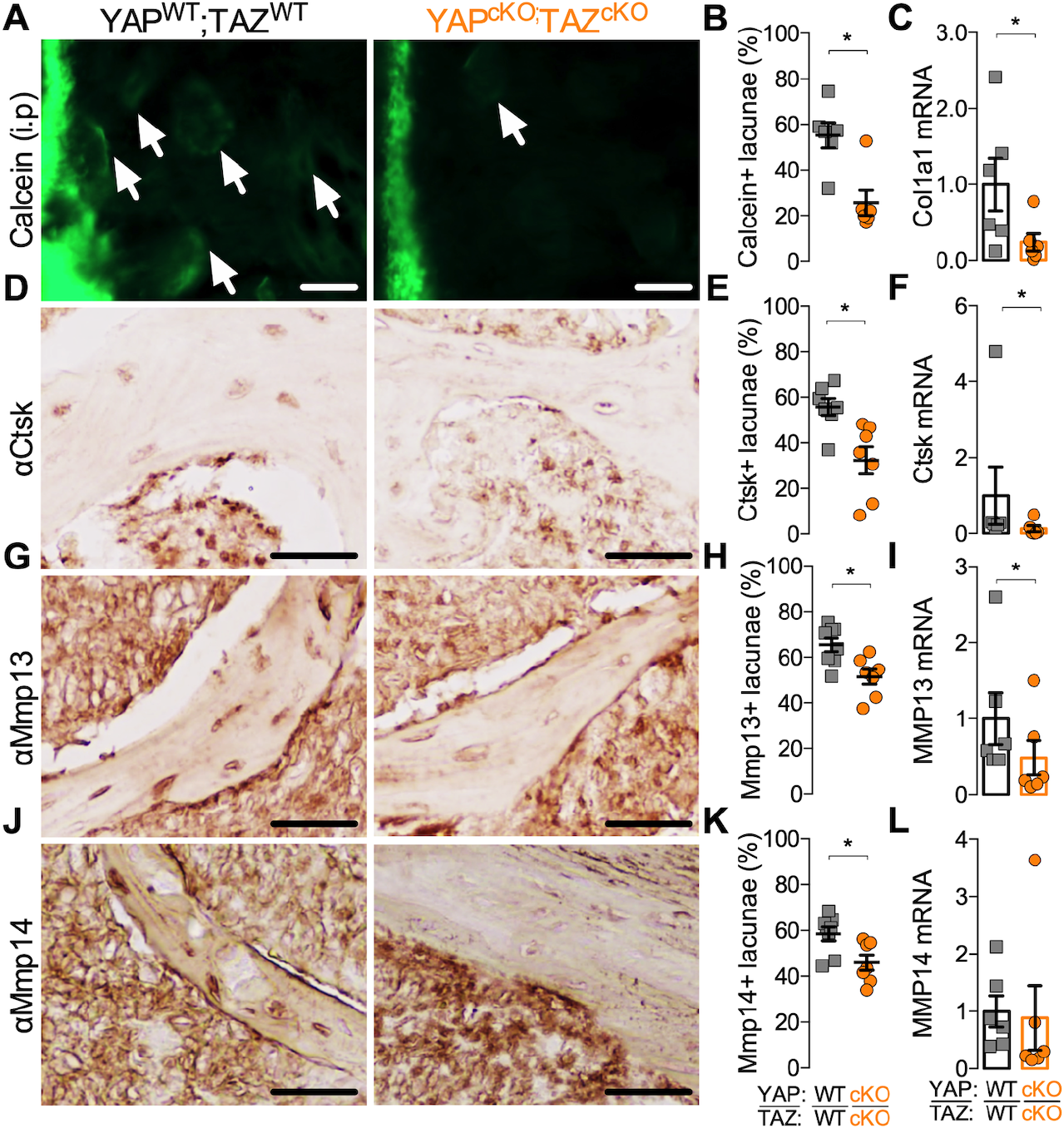
YAP/TAZ ablation from DMP1-expressing cells reduced osteocyte-mediated bone remodeling. **A)** Representative double fluorochrome-labeled images in cortical bone from P28 femurs (Calcein label injected at P21). **B-C)** Quantification of **(B)** Calcein positive lacunae and **(C)** relative transcript expression for collagen1a1 (Col1a1) in P84 femoral bone preparations. **D)** Representative micrographs of cancellous metaphyseal bone from P84 femurs immunostained for cathepsin K (Ctsk). **E-F)** Quantification of **(E)** Ctsk immunostained lacunae and **(F)** relative transcript expression for Ctsk in P84 femoral bone preparations. **G)** Representative micrographs of cancellous metaphyseal bone from P84 femurs immunostained for matrix metalloproteinase 13 (Mmp13). **E-F)** Quantification of **(E)** Mmp13 immunostained lacunae and **(F)** relative transcript expression for Mmp13 in P84 femoral bone preparations. **J)** Representative micrographs of cancellous metaphyseal bone from P84 femurs immunostained for matrix metalloproteinase 14 (Mmp14). **K-L)** Quantification of **(K)** Mmp14 immunostained lacunae and **(L)** relative transcript expression for Mmp14 in P84 femoral bone preparations. Data presented as individual samples with lines corresponding to the mean and standard error of the mean (SEM). Sample sizes, n = 6-8. Scale bars equal 30 μm in all images.

In addition to local matrix deposition, osteocytes also locally degrade their extracellular matrix. Many of the matrix metalloproteinases and other enzymes employed by osteoclasts to resorb bone are also expressed by osteocytes and are critical to perilacunar/canalicular remodeling (16, 41, 46). Therefore, to test whether YAP/TAZ regulate direct matrix remodeling by osteocytes, we measured mRNA and protein expression of the matrix proteases, cathepsin k (Ctsk), matrix metalloproteinase-13 (Mmp13), and matrix metalloproteinase-14 (Mmp14). Conditional YAP/TAZ deletion reduced the percentage of osteocytes that stained positive for Ctsk (Fig. 7D,E), Mmp13 (Fig. 7G,H), and Mmp14 (Fig. 7J,K) and decreased mRNA expression of *Ctsk* (Fig. 7F) and *Mmp13* (Fig. 7I), but not *Mmp14* (Fig. 7L) *in vivo*.

### YAP/TAZ inhibition abrogated TGFβ-induced remodeling gene expression in vitro

Though unexplored in osteocytes, YAP and TAZ are known to mediate TGF-β signaling in a variety of other cell types (30–33). Further, either pharmacologic inhibition or genetic ablation of TGF-β receptors from osteocytes caused defective perilacunar/canalicular remodeling and associated gene expression (29). Therefore, we next tested whether YAP/TAZ mediate TGF-β signaling, Cyr61/Ctgf expression, and PLR-associated gene expression in osteocytes by treating two osteocyte-like cell lines with combinatorial TGF-β and/or verteporfin (VP). VP is a small molecule inhibitor that physically blocks the interaction of YAP and TAZ with their transcriptional co-effectors, particularly the TEAD transcription factors, preventing YAP/TAZ-TEAD transcriptional activity (36, 47).

We first used mouse osteocyte-derived IDG-SW3 cells, which carry a Dmp1 cis-regulatory system driving GFP expression, as a marker of living osteocytes (48, 49). As shown previously, IDG-SW3 cells exhibited robust DMP1-GFP transgene expression after 21 days of *in vitro* osteocytic differentiation (Fig. 8A). Beginning at day 20, cells were treated with vehicle (DMSO) or 3 µM verteporfin (Fig. 8A). At day 21, cells were treated with 5 ng/ml TGF-βor PBS vehicle for 6 hours and gene expression was evaluated by q-PCR. Similar experiments were conducted in mouse osteocyte-derived OCY454 cells (50) (Fig. S7A).

**Figure 8.**
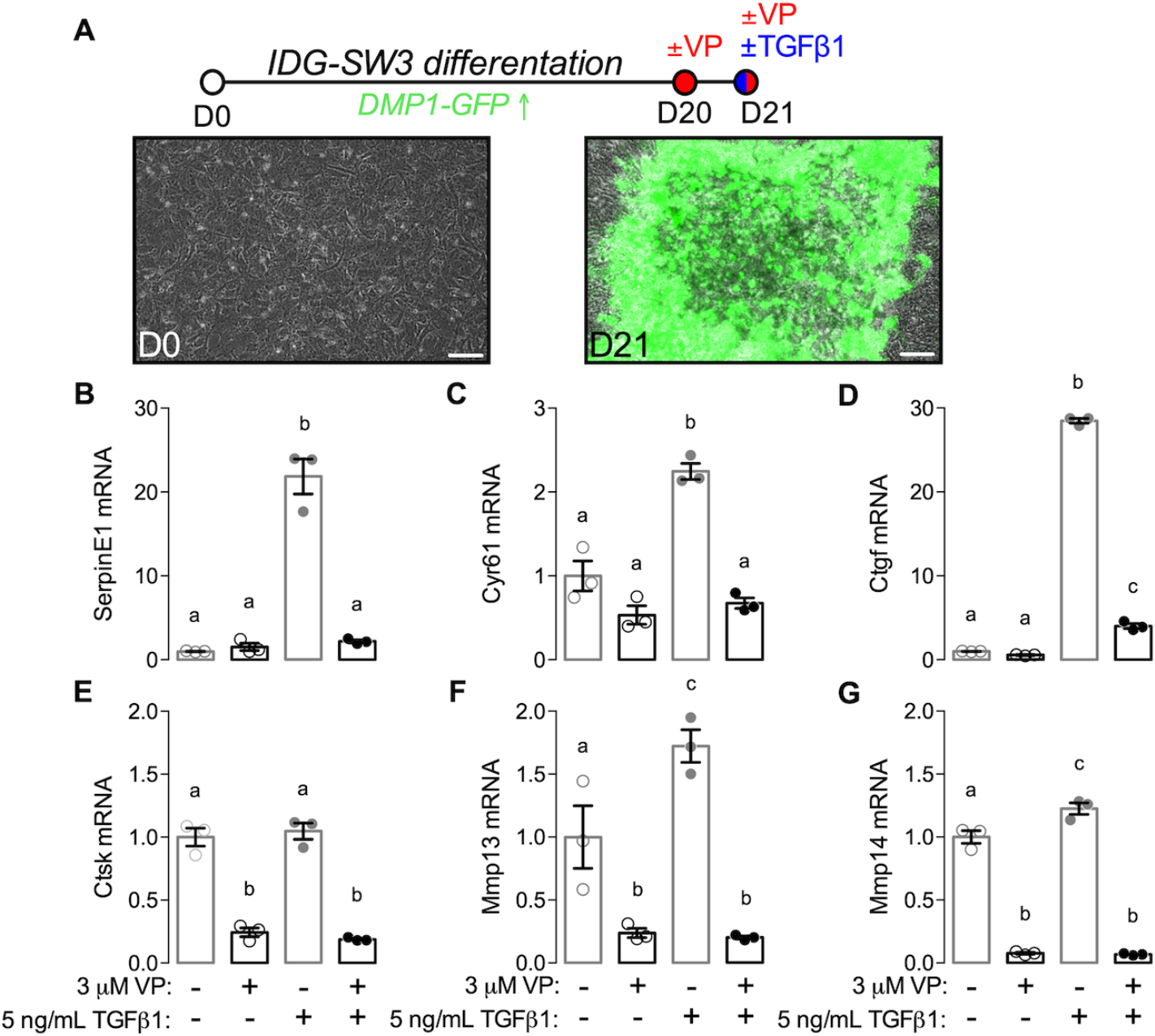
Inhibition of YAP/TAZ-TEAD with verteporfin (VP) reduced TGF-β-induced PLR gene expression *in vitro*. **A)** Osteocyte-like IDG-SW3 cells were differentiated for 21 days, with osteocyte differentiation reported by DMP1-GFP transgene expression. Cells were combinatorially treated with inhibitor verteporfin (VP) and/or 5 ng/ml TGFβ1 at day 21. **B-G)** mRNA expression, normalized to 18s rRNA, was evaluated for **(B)** serpin family E member 1 (SerpinE1), **(C)** cysteine-rich angiogenic inducer 61 (Cyr61), **(D)** connective tissue growth factor (Ctgf), **(E)** cathepsin K (Ctsk, **(F)** matrix metalloproteinase-13 (Mmp13), and **(G)** matrix metalloproteinase-14 (Mmp14). Relative expression was expressed as fold vs. vehicle (PBS + DMSO)-treated cells. Sample sizes, N = 3.

First, to test whether verteporfin blocked TGF-β-induced YAP/TAZ signaling, we evaluated expression of serpin family E member 1 (*SerpinE1)* and expression of known osteocyte-to-osteoblast/osteoclast paracrine signaling factors, *Cyr61* and *Ctgf*. Each of these genes are established TGF-β-inducible, YAP/TAZ-TEAD target genes (24, 25, 51, 52). As expected, TGF-βrobustly induced *SerpinE1*, *Cyr61*, and *Ctgf* mRNA expression, which was abrogated by verteporfin treatment (Fig. 8B-D). Next, to test whether TGF-β-induced YAP/TAZ activation regulated expression of genes associated with perilacunar/canalicular remodeling, we quantified mRNA expression of *Ctsk, Mmp13*, and *Mmp14*. TGF-β significantly induced expression of *Mmp13* and *Mmp14*, but not *Ctsk*, in IDG-SW3 cells, and verteporfin treatment abolished expression of all three genes (Fig. 8E-G). Similarly, in OCY454 cells, verteporfin abrogated TGF-βinduced expression of *Ctgf, Cyr61* and *SerpinE1* mRNA (Fig. S7B-D) as well as *Mmp14* and *Ctsk*, but not *Mmp13* (Fig. S7E-G).

## DISCUSSION

This study enhances our understanding of how bone remodeling contributes to skeletal fragility by identifying the role of key transcriptional regulators in direct and indirect osteocyte-mediated remodeling. Here, we show that YAP and TAZ act downstream of TGF-β signaling in osteocytes to control bone matrix accrual, organization, and mechanical properties by regulating both perilacunar/canalicular remodeling and osteoblast/osteoclast activity. Osteocyte-conditional YAP/TAZ deletion indirectly reduced bone mass by decreasing osteoblast number and activity and increasing osteoclast activity, but also impaired osteocyte-intrinsic pericellular matrix deposition and degradation, impairing canalicular network connectivity without altering lacunar morphology. Mechanistically, we found that YAP/TAZ activity was required for TGF-β1 induction of matricellular growth factors involved in paracrine signaling from osteocytes to osteoblasts/osteoclasts and matrix proteases necessary for perilacunar/canalicular remodeling. Together, these data identify the transcriptional co-activators YAP and TAZ as key mediators of bone remodeling.

The roles of YAP and TAZ in skeletal-lineage cells are beginning to beclarified. Previously, we reported that dual YAP/TAZ deletion from osteoprogenitor cells and their progeny using Osterix-Cre mimicked severe cases of osteogenesis imperfecta (36). We identified a combinatorial role for YAP and TAZ in promoting osteoblast number and activity and suppressing osteoclast activity (36). Similarly, YAP deletion later in the osteoblast lineage, from committed osteoblasts using Osteocalcin-Cre, significantly reduced bone formation, further supporting a role for YAP in promoting osteogenesis *in vivo* (53). That study did not observe changes in osteoclast activity, potentially due to compensatory effects of TAZ or the Cre line used (53). We show here that 8kb-DMP1-Cre-conditional YAP/TAZ deletion from osteocytes also reduced osteoblast number and activity and increased osteoclast activity. Consistent with our data (40), Xiong et al. found that YAP/TAZ deletion using 10kb-DMP1-Cre reduced bone formation and increased osteoclast numbers (54). Notably, neither study observed significant changes in transcript expression of *SOST*, or *Opg/Rankl*, other factors known to mediate osteocyte signaling to osteoblast and osteoclasts (55).

Here, we found that YAP/TAZ deletion *in vivo* and inhibition *in vitro* reduced transcript expression of the matricellular growth factors, Cyr61 and Ctgf. Both Cyr61 and Ctgf are induced by TGF-β(51, 52) and are transcriptionally regulated directly by the YAP/TAZ-TEAD complex (24, 25). Cyr61 and Ctgf are expressed by osteocytes (17) and have been implicated in both osteoblastogenesis (21, 22) and osteoclastogenesis (23, 56). During osteoblastogenesis *in vitro*, Cyr61 enhances mesenchymal stem cell migration and regulates Wnt3A-induced osteogenic differentiation (21). Ctgf also enhances osteoblastogenesis *in vitro*, in part by inhibiting Notch signaling and inducing HES-1 transcription and NFAT transactivation (22)*. In vivo* deletion of Ctgf in Osteocalcin-expressing cells led to a mild low bone mass phenotype in male mice without alterations in osteoblast or osteoclast numbers (57). Similarly, deletion of Cyr61 at various stages of the osteoblast lineage (Osterix-Cre, Collagen1(2.3kb)-Cre, and Osteocalcin-Cre) resulted in low bone mass phenotypes of similar severity, suggesting mature osteoblasts/osteocytes are the primary source of Cyr61 in bone (18). Deletion of Cyr61 from osteocalcin-expressing mature osteoblasts also increased osteoclast numbers *in vivo* (18), consistent with the effects of osteocyte-conditional YAP/TAZ deletion here. Cyr61 inhibits osteoclastogenesis *in vitro* through a RANKL-independent mechanism (23), while Ctgf has been observed to promote osteoclast-precursor fusion through interaction with dendritic cell-specific transmembrane protein (DC-STAMP) (56). Future studies will be required to dissect the varied mechanisms by which osteocytes direct osteoblast/osteoclast-mediated remodeling, but these data identify a role for YAP/TAZ in indirect osteocyte-mediated bone remodeling by promoting osteoblast activity and suppressing osteoclast activity and implicate a TGF-β–YAP/TAZ-TEAD signaling axis in Cyr61 and Ctgf expression by osteocytes.

In addition to regulating indirect osteocyte-mediated coordination of osteoblasts and osteoclasts, we found that YAP/TAZ deletion also impaired perilacunar/canalicular remodeling of the bone matrix directly by osteocytes. TGF-β signaling is an osteocyte-intrinsic regulator of perilacunar/canalicular remodeling (29). Here, we found that DMP1-conditional YAP/TAZ deletion phenocopied the effects of both DMP1-conditional TGF-βreceptor deletion and pharmacologic inhibition, which impaired canalicular network length and reduced expression of matrix-degrading enzymes required for perilacunar/canalicular remodeling (i.e. Mmp13, Mmp14, and Ctsk) without altering lacunar morphology (29), as observed here. Confirming the function of these regulated genes, DMP1-conditional YAP/TAZ deletion also partially phenocopied knockout mouse models of these important matrix proteases. First, global knockout of Mmp13 in mice impaired osteocyte perilacunar/canalicular remodeling, reducing flourochrome-labeled mineral deposition around osteocytes and canalicular network connectivity (41). Second, global knockout of Mmp14 in mice significantly reduced canalicular network development and maintenance, reducing osteocyte processes density and length (46). Lastly, global knockout of Ctsk in mice impaired bone matrix collagen organization that increased bone fragility in spite of high bone mass and increased osteoclast activity (58). The concordance of these observations led us to hypothesize that YAP/TAZ act downstream of TGF-βto regulate perilacunar/canalicular remodeling genes in osteocytes.

TGF-β signaling is known to regulate YAP/TAZ in a variety of cell types (30–33). For example, in LM2-4 metastatic breast cancer cells, siRNA-depletion of YAP/TAZ and/or TEAD reduced expression of *Ctsk, Mmp13*, and *Mmp14* after 24 hours of treatment with TGF-β(59). Furthermore, sh-RNA interference of YAP1 in pancreatic ductal adenocarcinoma cells downregulated Mmp13 protein expression (60). In human fibroblasts, downstream induction of Mmp13 by TGF-βdepends on the activation of p38α and Smad3 (61). These observations parallel our findings in osteocytes.

The mechanisms by which TGF-βregulate YAP and TAZ continue to emerge. YAP/TAZ form complexes with the R-Smad proteins and co-activate Smad transcriptional activity to regulate Smad-target gene expression (32, 33). Furthermore, dual YAP/TAZ inhibition activates the transcription factor AP-1 to induce Smad7-mediated inhibition of R-Smad complex formation and suppress TGF-β/Smad target gene expression (30). In a distinct Smad3-independent mechanism, TGF-βinduced TAZ transcriptional activity through a MRTF-mediated, p38- and redox-dependent manner (31). Together these data position YAP and TAZ as potential downstream mediators of TGF-β-induced bone remodeling gene expression in osteocytes. Ongoing work, beyond the scope of the present study, will identify the transcriptional mechanisms by which YAP/TAZ mediate TGF-βinduction of perilacunar/canalicular remodeling genes in osteocytes.

Perilacunar/canalicular remodeling and osteoblast/osteoclast-mediated remodeling coordinately determine skeletal strength (5, 6). Compromised bone strength caused by defects in bone mass and/or quality results in fragility fracture (4–6). Bone quantity is primarily regulated by the balance in of bone turnover rate while quality includes matrix composition and microarchitectural geometry (4, 5). Both of these factors contribute to the pathogenesis of skeletal fragility diseases such as osteoporosis, Paget’s disease, and osteogenesis imperfecta (5). Here, we found that osteocyte-conditional YAP/TAZ deletion decreased bone mass and altered collagen matrix content and organization, producing moderate defects in bone mechanical behavior. Osteocyte-specific knockouts did not exhibit the spontaneous bone fractures prevalent in mice lacking YAP/TAZ in the full osteoblast lineage (36), but their defects correspond with the increased bone fragility described in other models of defective perilacunar/canalicular remodeling (29, 41). Age-related decreases in canalicular network connectivity are associated with microdamage accumulation that contributes to age-associated skeletal fragility (62), further suggesting a contribution of the canalicular network to bone strength.

Osteocytes are the key mechanosensing cells in bone (63), and YAP and TAZ are key mediators of mechanotransduction in many cell types (64), but the putative roles of YAP and TAZ in osteocyte mechanotransduction are unknown. In addition, as mechanical loading also induces perilacunar/canalicular remodeling (65), continued investigation is warranted to determine the extent to which YAP and TAZ mediate load-induced bone adaptation and perilacunar/canalicular remodeling.

In conclusion, this study identifies the transcriptional co-activators, YAP and TAZ, as TGF-β-inducible regulators of bone quantity and quality that mediate osteocyte regulation of osteoblast and osteoclast activity and perilacunar/canalicular remodeling. Specifically, osteocyte YAP and TAZ are required for TGF-β-induced expression of paracrine growth factors critical to the formation of osteoblasts/osteoclasts (i.e., Cyr61, Ctgf) and of enzymes that affect perilacunar/canalicular remodeling. Further elucidation of the mechanisms by which the TGF-β and YAP/TAZ signaling pathways regulate osteocyte function may help guide targeted therapeutic strategies for the treatment of skeletal fragility diseases.

## MATERIALS AND METHODS

### Animals

Mice harboring loxP-flanked exon 3 alleles in both YAP and TAZ were kindly provided by Dr. Eric Olson (University of Texas Southwestern Medical Center). All protocols were approved by the Institutional Animal Care and Use Committees at the University of Notre Dame and the University of Pennsylvania and in adherence to federal guidelines for animal care. Osteocytes were targeted using Cre-recombination under the control of an 8kb fragment of the dentin matrix protein-1 promoter (8kb-DMP1-Cre) (35). Mice with homozygous floxed alleles for both YAP and TAZ (YAP^fl/fl^;TAZ^fl/fl^) were mated with double heterozygous conditional knockout mice (YAP^fl/+^;TAZ^fl/+^;DMP1-Cre) to produce eight possible genotypes in each litter, but only the mice in Table 1 were compared. Both male and female mice were evaluated with YAP^fl/fl^;TAZ^fl/fl^ mice serving as littermate wild type (WT) controls. All mice were fed regular chow ad libitum and housed in cages containing 2-5 animals each. Mice were maintained at constant 25°C on a 12-hour light/dark cycle. Mice were tail or ear clipped after weaning or prior to euthanasia and genotyped by an external service (Transnetyx, Inc.).

### Histology and histomorphometric analysis

Femurs and tibias from P28 and P84 mice were fixed with 10% neutral buffered formalin for 48 hours at 4°C and decalcified for 4 weeks with 0.25M EDTA (pH 7.4) at 4°C. Paraffin sections (5 μm thickness) for were processed for either immunohistochemistry or histology. Primary antibodies were compared to both normal rabbit sera and no primary control sections. For immunostaining, anti-Ctsk (1:75, ab19027; abcam), anti-Mmp13 (1:100, ab39012; abcam), anti-Mmp14 (1:100, ab38971; abcam), anti-YAP (1:200, 14074; Cell Signaling), and anti-TAZ (1:200 NB110-58359; Novus Biologicals) primary antibodies were applied overnight. This was followed by incubation with corresponding biotinylated secondary antibody, avidin-conjugated peroxidase, and diaminobenzidine substrate chromogen system (329ANK-60; Innovex Biosciences), which allowed for immunohistochemical detection of positively stained cells. To determine the number of positively immunostained cells, 3 fields of view per mouse per antibody were de-identified and manually scored as either positive or negative and reported as a percent positively stained per total number of osteocytes scored. Hematoxylin and eosin stains (H&E), Tartrate-resistant acid phosphatase (TRAP), Picrosirius Red, and silver nitrate stains were used to stain for osteoblasts, osteoclasts, collagen, and the osteocyte lacunar/canalicular network as previously shown (42, 43). Osteoblast number per bone surface, osteoclast surface per bone surface and osteocyte number per bone area were quantified using Osteomeasure^TM^ (OsteoMetrics) on H&E and TRAP stained sections, respectively. To determine canalicular length, primary canaliculi emanating from each lacuna and extending as a single, unbranched process were traced with ImageJ. The mean length was taken from at least 5 osteocytes per field, with 3 fields per mouse and 8 mice per group (29).

Cryohistology of mineralized tissue was performed as previously described (66). Briefly, P28 femurs were fixed with 10% neutral buffered formalin for 48 hours at 4°C, transferred to 30% sucrose in PBS overnight at 4°C, and then embedded in O.C.T. compound (Tissue-Tek). All sections (7 μm thickness) were made from undecalcified femurs using cryofilm IIC tape (Section Lab Co. Ltd.), which maintains morphology of mineralized sections. The taped sections were glued to microscope slides using UV adhesive glue. Taped sections were rehydrated and decalcified with 0.25M EDTA (pH 7.4) for 5 minutes prior to staining. Terminal deoxynucleotidyl transferase dUTP nick end labeling (TUNEL) assays were then performed using the Click-iT Plus TUNEL Assay kit (C10618; Invitrogen) according to the manufacturer’s instructions. On thick sections (20 μm thickness), F-actin was stained with Alexa Fluor 488-conjugated phalloidin (1:50, Life Technologies) for confocal imaging.

Methyl-methacrylate-embedded bones from mice injected with Calcein (C0875-25G; Sigma-Aldrich) and Alizarin Complexone (A3882-25G; Sigma Aldrich) were processed for dynamic bone histomorphometry. Using a diamond-embedded wire saw (Delaware Diamond Knives), transverse sections (40 µm) were cut from the midshaft and ground to a final thickness of 20 µm. The slice sections were mounted on slides, and three sections per limb were analyzed using Osteomeasure^TM^ (OsteoMetrics). The following primary data were collected: total bone surface length (BS); single label perimeter (sL.Pm); double label perimeter (dL.Pm); and double label width (dL.Ith). From primary data, we derived mineralizing surface (MS/BS = [1/2sL.Pm+dL.Pm]/B.Pm ×100; %); mineral apposition rate (MAR = dL.Ith/5 days; µm/day) and bone formation rate (BFR/BS = MAR × MS/BS; µm^3^/µm^2^ per day). Six mice were analyzed per group.

### Microcomputed tomography

Harvested femurs from P84 mice were stored at − 20°C until evaluation. Frozen specimens were thawed and imaged using a vivaCT 80 scanner (Scanco Medical) to determine trabecular and cortical femoral bone architecture prior to mechanical testing. The mid-diaphysis and distal femur were imaged with an X-ray intensity of 114 μA, energy of 70 kVp, integration time of 300 ms, and resolution of 10 μm. Mid-diaphyseal and distal femoral 2D tomograms were manually contoured, stacked and binarized by applying a Gaussian filter (sigma =1, support =1) at a threshold value of 250 mg HA/cm^3^. Eight mice were analyzed per group.

### Mechanical testing

Three-point bend testing to failure was carried out on femurs from 12-week-old mice. The femurs were loaded with the condyles facing down onto the bending fixtures with a support span length of 4.4 mm. The upper fixture was aligned with the mid-diaphysis. The femurs were loaded to failure at a rate of 0.5 mm/s using the ElectroForce 3220 Series testing system (TA Instruments). Stiffness, maximum load to failure, work to maximum load and work to failure were quantified using a custom MATLAB script (36). Eight mice were analyzed per group.

### Imaging

Histological and immunohistochemical sections were imaged on either a 90i Upright/Widefield Research Microscope (Nikon Instruments) at the 20× and 40× objectives or an Axio Observer Z1 (Zeiss) at the 20×, 40×, and 63× objectives. The samples tested in three-point bending were stained with Picro-sirius Red to construct a multivariate regression model that included collagen organization and content as a predictor of mechanical behavior. These samples were imaged both under polarized light using an Eclipse ME600 Microscope (Nikon Instruments) at the 20× objective and using second harmonic generated (SHG) microscopy. SHG images were taken on a multiphoton-enabled Fluoview Research Microscope (Olympus) at a fundamental wavelength of 875nm with the 25× objective on sections oriented in the same direction for all groups. All SHG images were quantified using ImageJ and reported as mean pixel intensity within the cortical region relative to WT bone. Mean pixel intensities across four separate regions of interest within each image of the cortex were averaged as technical replicates for a given histological section. Confocal images were acquired using an LSM 710 confocal (Zeiss) with the 63× objective in 20-micron thick sections. Three separate regions of interest within the cortex were averaged as technical replicates for a given histological section. All confocal image stacks were quantified using ImageJ. Briefly, image stacks were smoothed by applying a Gaussian filter (sigma = 0.3), then skeletonized using the Skeletonize3D function. Finally, the AnalyzeSkeleton2D/3D function was used to quantify mean branch length, number of branches, and number of junctions (67).

### Osteocyte lacunae visualization and quantification

Nano-computed tomography (nanoCT) was used toevaluate the morphological characteristics of osteocyte lacunae in the proximal tibia (Xradia Versa XRM-520, Zeiss, Dublin, CA). Tibiae were harvested and removed of all non-osseous tissue. Samples were scanned in 70% EtOH, in custom-made sample holders that oriented the samples vertically on the stage. Tibias were originally scanned with a 4× objective to determine regions of interest. A 650 µm^3^ region of bone was located on the anterior medial aspect of the tibia, 8 mm from the tibiofibular junction (TFJ). Nano-computed tomography images image were collected in the region of interest using a 20× objective with energy settings of 40 V, 3.0 W, using the Air Filter and 3,201 projections. To obtain a constant resolution of 0.6 µm voxel for all the samples, the source and detector distance were varied between 5.8 and 8.5 mm from the sample, and excitation ranged from 6 − 8 seconds contingent on resulting intensity values.

Dragonfly 3.6 (Object Research Systems (ORS) Inc, Montreal, QC) was used for segmentation of bone types and osteocyte lacunae regions. Cortical and trabecular bone were manually segmented, while osteocyte lacunae were identified and segmented in three dimensions using a global threshold for each sample. Segmented regions based on global thresholding were further filtered to more accurately capture the potential osteocyte lacunae distribution. Objects that were outside of previously reported values (< 50 *µ*m^3^ and > 1,500 *µ*m^3^) (68–70) to be an osteocyte lacuna were removed. Using Dragonfly volume, surface area, aspect ratio, phi, and theta were measured for each osteocyte. Additionally, custom python code based previous work (68, 71, 72) was implemented to analyze osteocyte lacunar shape measures such as oblateness, stretch, primary orientation, secondary orientation, height, width, length, and were also computed for each osteocyte lacunae. Oblateness and stretch are calculated from the eigenvalues of the shape tensor for each individual osteocyte lacunae and are defined previously (68).

### Cell culture

OCY454 cells were cultured in α-minimum essential media (α-MEM) supplemented with 10% fetal bovine serum and 1% antibiotic/antimycotic (Gibco) at 33°C. IDG-SW3 cells were cultured in α-MEM supplemented with 10% fetal bovine serum, 1% antibiotic/antimycotic (15240062; Gibco) and 1 U/mL INF-γ (PMC4031; Invitrogen) at 33°C. Prior to treatment, OCY454 cells were differentiated for 12 days at 37°C while IDG-SW# cells were differentiated for 21 days at 37°C in α-MEM supplemented with 10% fetal bovine serum (100-106; Gemini Bio-Products), 1% antibiotic/antimycotic (15240062; Gibco), 50 μg/ml ascorbic acid (A-4544; Sigma-Aldrich), and 4 mM β-glycerophosphate (G-9422; Sigma-Aldrich) without INF-γ. Prior to treatment on the last day of differentiation, both OCY454 and IDG-SW3 cells were treated with 3 μM verteporfin (SML0534; Sigma-Aldrich) overnight in 1% fetal bovine serum. The next morning, the media was changed and supplemented with 3 μM verteporfin and 5ng/ml TGF-β1 (HZ1011; Humanzyme) for 6 hours before mRNA isolation.

### RNA isolation and qPCR

Femur samples were dissected, the ends were cut and discarded, and marrow flushed before being snap-frozen in liquid nitrogen-cooled isopentane for 1 minute prior to storage at −80°C until processing. Tissue was then homogenized via mortar and pestle and RNA from the sample was collected using Trizol Reagant (15596026; Life Technologies) followed by centrifugation in chloroform. RNA from femur tissue and cell culture experiments were purified using the RNA Easy Kit (74106; Qiagen) and quantified by spectrophotometry using a NanoDrop 2000 (Thermo-Fisher Scientific).

Reverse transcriptase polymerase chain reaction (RT-PCR) was performed on 0.5 μg/μl concentration of RNA using the High-Capacity cDNA Reverse Transcription Kit (4368814; Thermo-Fisher Scientific). Quantitative polymerase chain reaction (qPCR) assessed RNA amount using a StepOnePlus^TM^ Real-Time PCR System (Thermo-Fisher Scientific) relative to the internal control of 18s ribosomal RNA (18s rRNA). Data are presented using the ΔΔCt method. Eight mice per group were used. Specific mouse primer sequences are listed (Table 2).

**Table 2:**
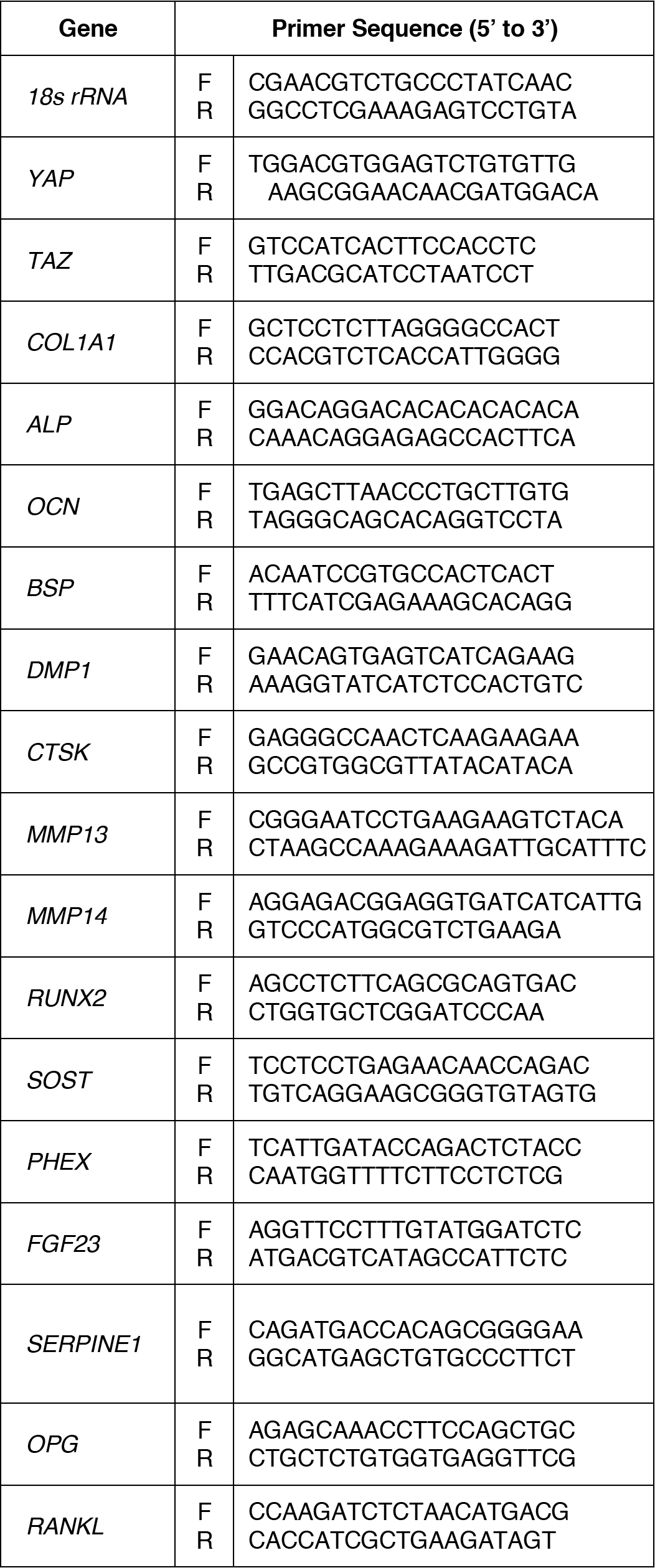
qPCR primers. Mouse primers used for qPCR

### Statistics and regression

Sample sizes were selected *a priori* by power analysis based on effect sizes and population standard deviations taken from published data on YAP^fl/fl^;TAZ^fl/fl^ mice in other tissues (73), assuming a power of 80% and α=0.05. All statistics and regression analyses were performed in GraphPad Prism or using R (Version 2.13.1). Comparisons between two groups were made using the two-tailed student’s t-test, provided the data were normally distributed according to D’Agostino-Pearson omnibus normality test and homoscedastic according to Bartlett’s test. When parametric test assumptions were not met, data were log-transformed, and residuals were evaluated. If necessary, the non-parametric Mann-Whitney test was used. A p-value < 0.05 was considered significant. Data are represented as individual samples with mean ± standard error of the mean (s.e.m.). All statistics and regression analyses for osteocyte lacunae quantification were performed using R version 3.5.1. A repeated-measures ANOVA with post hoc Tukey’s HSD comparison test was used to analyze the osteocyte lacunae morphology parameters. Multivariate analysis was performed as described previously (74), with some modifications (36). Briefly, we used an exhaustive best subsets algorithm to determine the best predictors of maximum load and stiffness from a subset of morphological parameters measured, which included moment of inertia (I) or section modulus (I/c), tissue mineral density (TMD), and second harmonic generated (SHG) intensity based on the Akaike’s information criterion (AIC) (75) The lowest AIC selects the optimal model while giving preference to less complex models (those with fewer explanatory parameters). Finally, the overall “best” model for each predicted mechanical property was compared to the prediction from only the moment of inertia (I/c or I for maximum load and stiffness, respectively) using Type II general linear regression.

## ACKNOWLEDGEMENTS

YAP^fl/fl^;TAZ^fl/fl^ mice were provided by Dr. Eric Olson (University of Texas Southwestern Medical Center). Theresa Sikorski (University of Notre Dame) provided initial mouse husbandry and maintenance. The authors declare no conflicts of interest. This work was supported by the National Institute of Arthritis, Musculoskeletal, and Skin Diseases (NIAMS) of the National Institutes of Health through grants T32 AR007132 (to C.D.K.), P30 AR069619 (to J.D.B.), and R21 AR071559 (to J.D.B., A.G.R., and T.M.B.).

## AUTHOR CONTRIBUTIONS

C.D.K. and J.D.B. designed research; C.D.K., J.C.C., K.M.J., A.G.R., V.L.F., T.M.B., and J.D.B. analyzed data; C.D.K., J.C.C., K.M.J., and D.J.H. performed research; J.C.C., D.J.H., A.G.R., L.Q., V.L.F., and T.M.B. contributed new reagents/analytic tools; C.D.K. and J.D.B. wrote the paper; All authors reviewed and approved the paper.

---

**Supplemental Figure 1.**
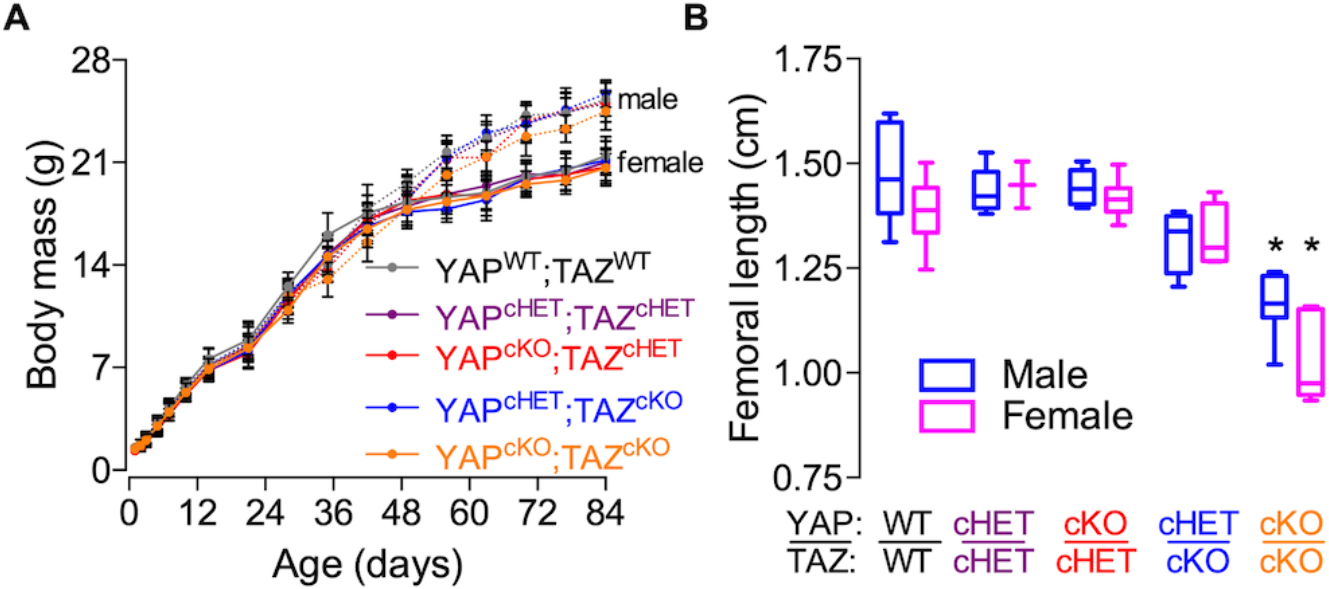
Combinatorial YAP/TAZ ablation from DMP1-expressing cells is functionally redundant. **A)** Body masses from mice with allele-dose dependent YAP/TAZ ablation from DMP1-expressing cells until P84. **B)** Femoral lengths from mice with allele-dose dependent YAP/TAZ ablation from DMP1-expressing cells at P84 for both males and females. Body mass data presented as mean with lines corresponding to the standard error of the mean (SEM). Sample sizes, N = 6-22 per sex per group, Femoral length data presented as a box and whiskers plots with whiskers corresponding to maximum and minimum values. Sample sizes, N = 2-10 per sex per group.

**Supplemental Figure 2.**
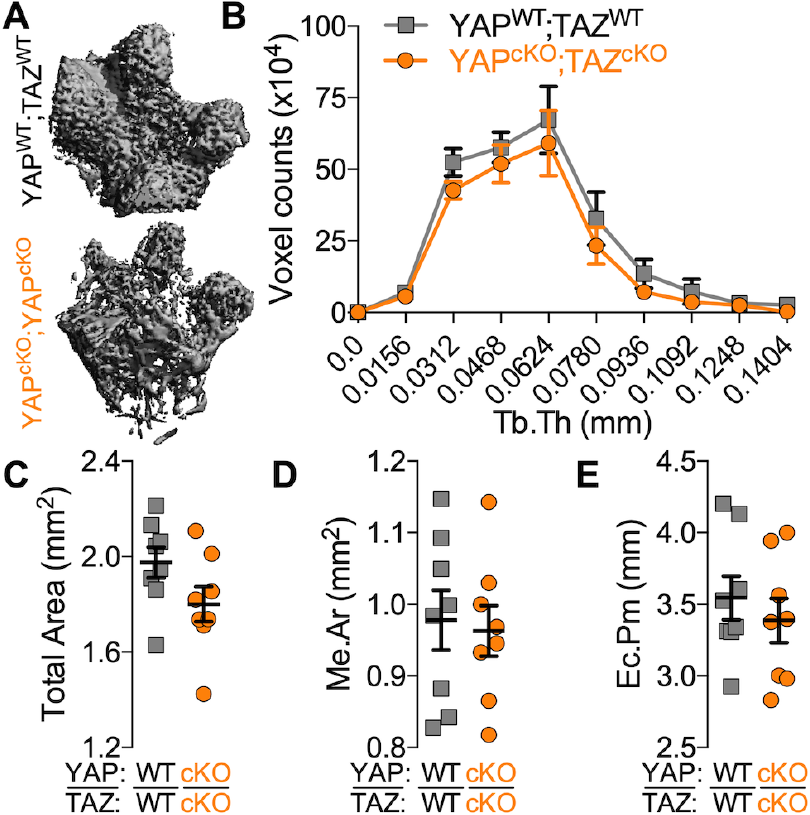
YAP/TAZ ablation from DMP1-expressing cells altered bone microarchitecture. **A)** Representative microCT reconstructions of distal metaphyseal microarchitecture in P84 femurs. **B)** Quantification of trabecular thickness distributions in cancellous bone. **C-E)** Quantification of cortical microarchitectural properties: **(C)** total area, **(D)** medullary area (Me.Ar), and **(E)** endocortical perimeter (Ec.Pm). Data are presented as individual samples with lines corresponding to the mean and standard error of the mean (SEM). Sample sizes, N = 8 per group. These data were originally presented in (40) but removed from (36).

**Supplemental Figure 3.**
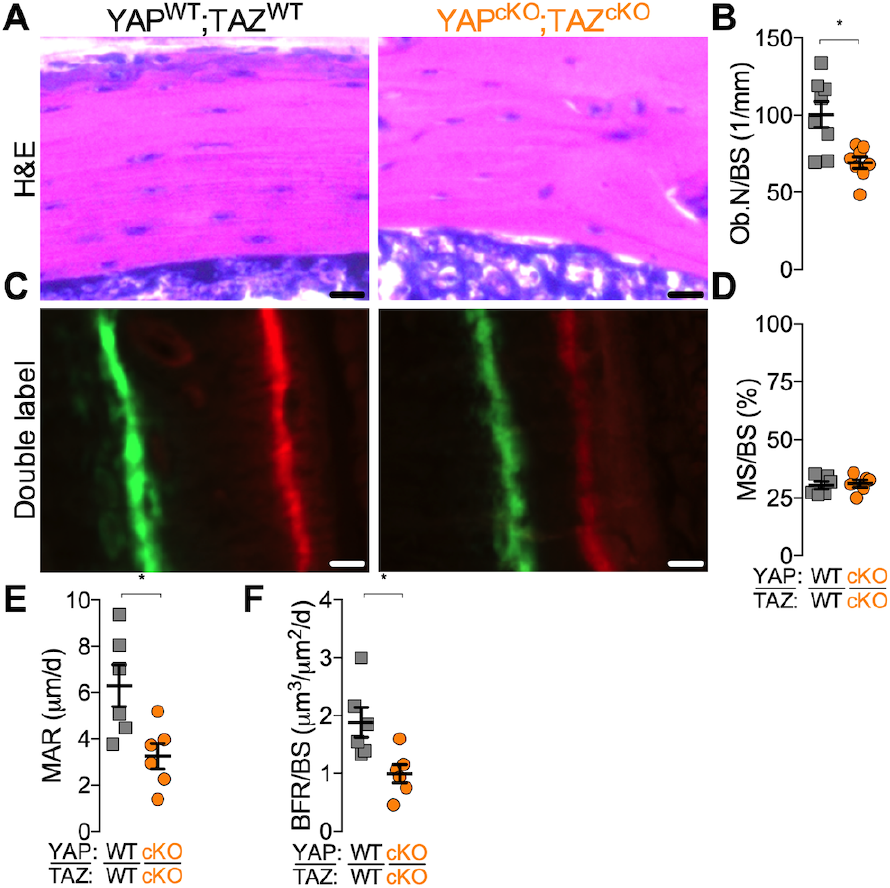
YAP/TAZ ablation from DMP1-expressing cells altered cortical osteoblast number and function. **A)** Representative micrographs of P84 cancellous metaphyseal bone stained by H+E. **B)** Quantification of osteoblast number per bone surface (Ob.N/BS). **C)** Representative micrographs of double flourochrome labeled cortical bone in P28 femurs. **D-F)** Quantification of **(D)** mineralizing surface percentage (MS/BS), **(E)** mineral apposition rate (MAR) and **(F)** bone formation rate (BFR/BS). Data presented as individual samples with lines corresponding to the mean and standard error of the mean (SEM). Sample sizes N = 6-8. Scale bars indicate 30 μm.

**Supplemental Figure 4.**
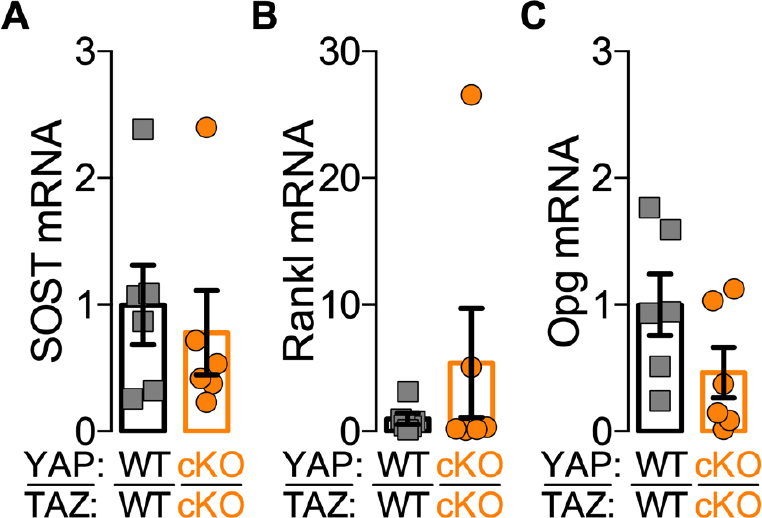
YAP/TAZ ablation from DMP1-expressing cells did not affect canonical markers of indirect osteocyte-mediated bone remodeling. **A-C)** Femoral bone preparations from P84 mice were harvested to quantify mRNA expression. Expression levels, normalized to 18S rRNA, were evaluated for **(A)** sclerostin (SOST), **(B)** receptor activator of nuclear factor kappa-B ligand (Rankl), and **(C)** osteoprotegerin (Opg), Data presented as individual samples with lines corresponding to the mean and standard error of the mean (SEM). Sample sizes, N = 6 per group.

**Supplemental Figure 5.**
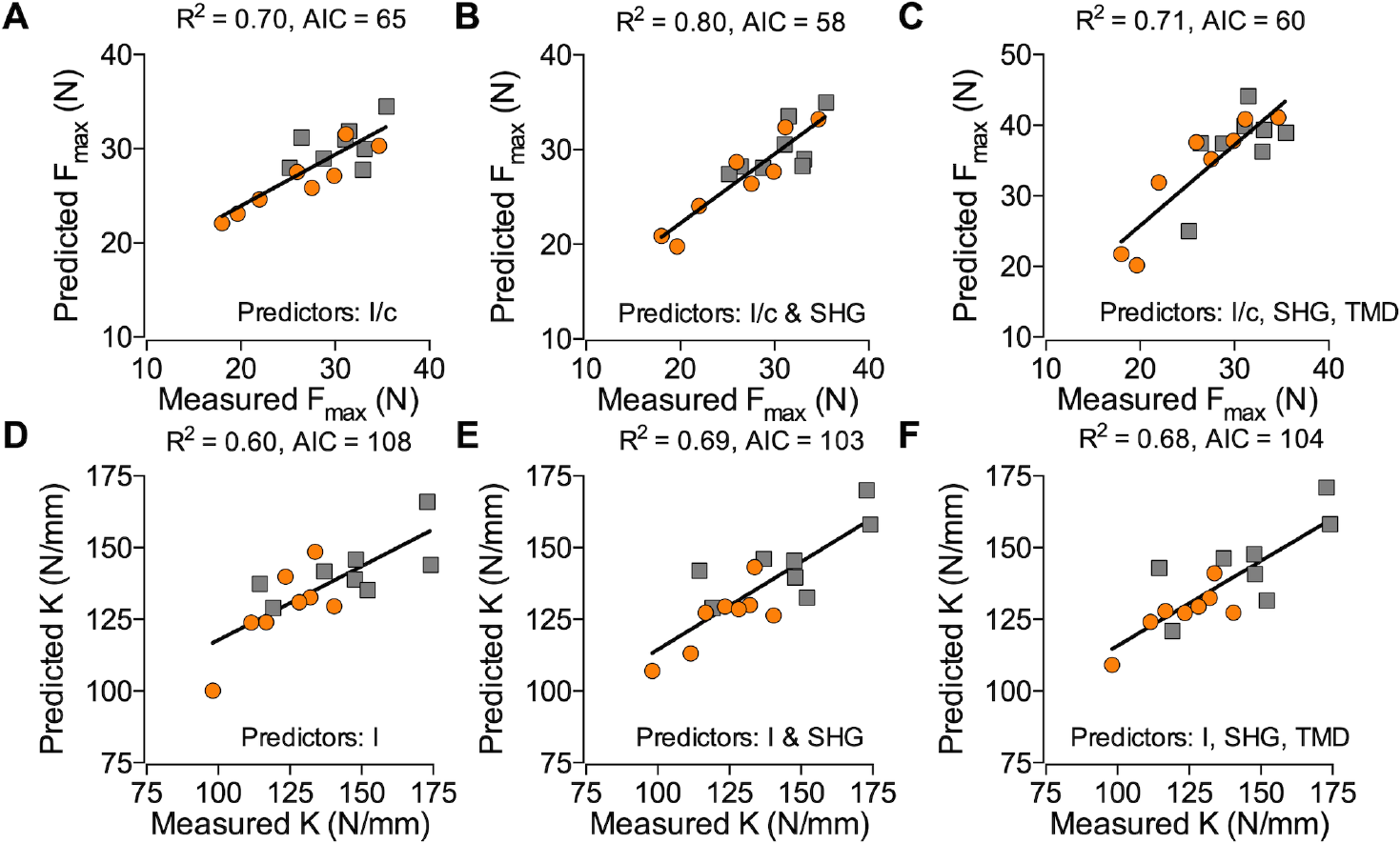
Best subsets analysis on morphological parameters from YAP/TAZ ablation in DMP1-expressing cells demonstrated increased goodness of fit parameters by accounting for collagen content and organization. Using the femurs that were scanned by microCT, tested in three-point bending, and imaged using SHG, experimental ultimate load (F_max_) values were compared with predicted model values with (**A**) only section modulus (I/c) as a predictor, (**B**) I/c and second harmonic generated signal intensity (SHG), or (**C**) I/c, SHG, and tissue mineral density (TMD). Experimental bending stiffnesses (K) were compared to predicted stiffnesses with (**D**) only moment of inertia (I) as a predictor, (**B**) I and SHG, or (**C**) I, SHG, and TMD. Akaike’s information criterion (AIC) and R^2^ values were used to analyze each model’s predictive power. Sample sizes, N = 8 per group. These data were originally presented in (40) but removed from (36).

**Supplemental Figure 6.**
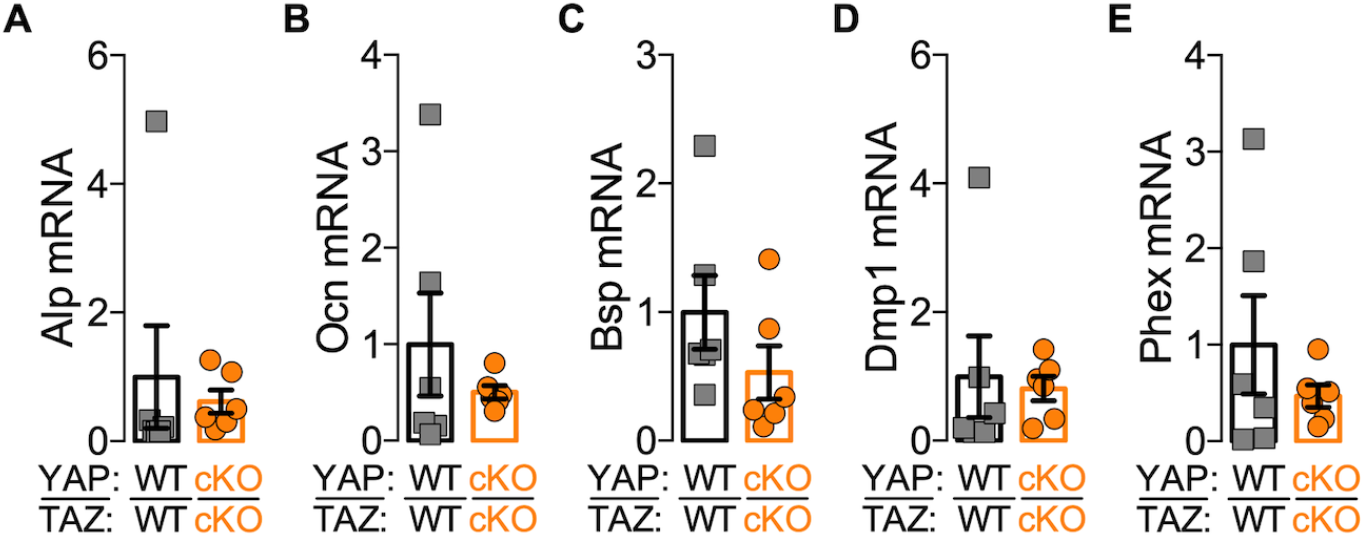
YAP/TAZ deletion did not alter osteogenic or osteocyte-marker gene expression. **A-E)** Femoral bone preparations from P84 mice were harvested to quantify mRNA expression. Expression levels, normalized to 18S rRNA, were evaluated for **(A)** alkaline phosphatase (Alp), **(B)** osteocalcin (Ocn)**, (C)** bone sialoprotein (Bsp), **(D)** dentin matrix protein 1 (Dmp1), and **(E)** phosphate-regulating neutral endopeptidase. Data presented as individual samples with lines corresponding to the mean and standard error of the mean (SEM). Sample sizes, N = 6 per group.

**Supplemental Figure 7.**
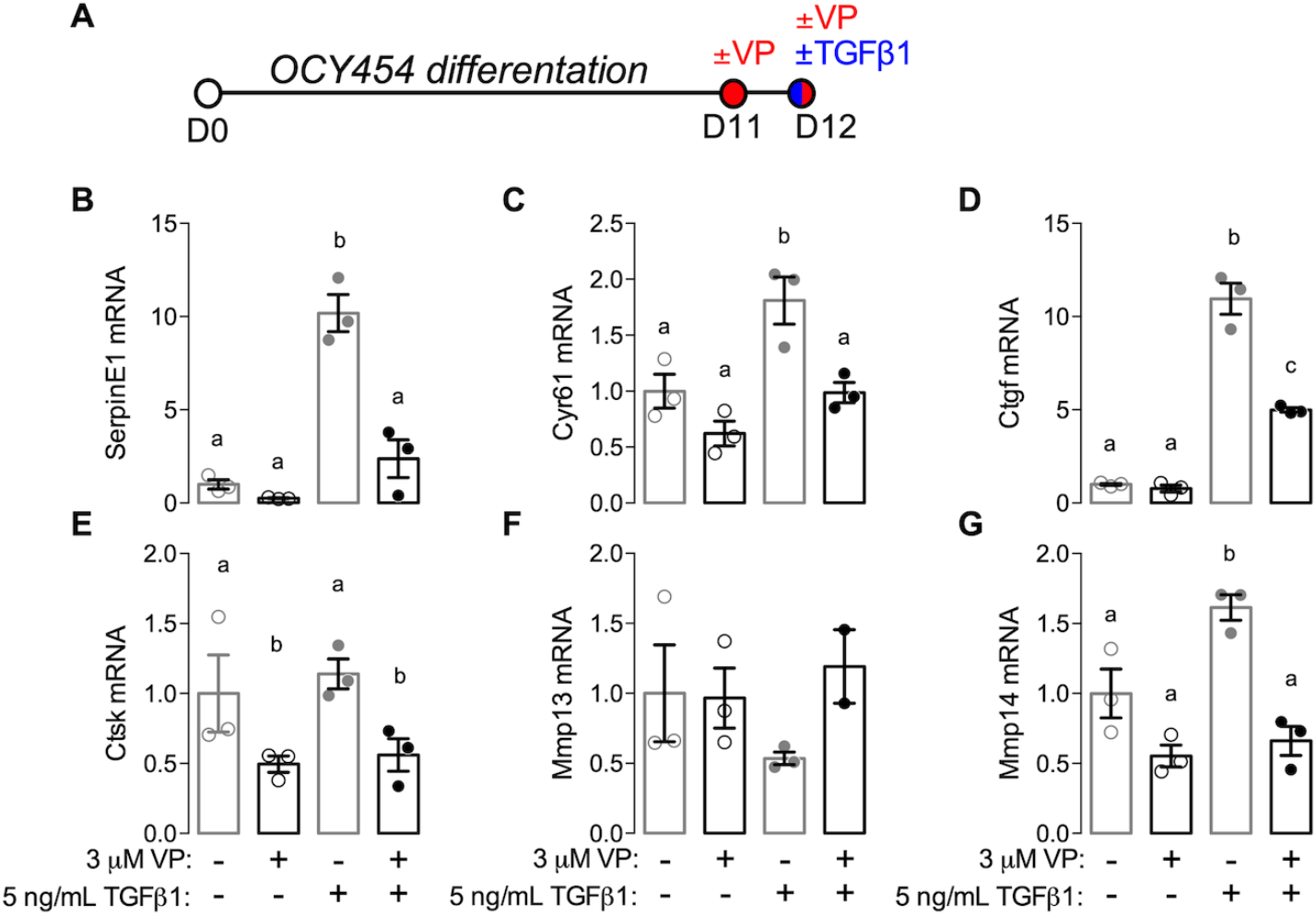
Inhibition of YAP/TAZ-TEAD with verteporfin (VP) reduced TGF-β-induced remodeling gene expression in OCY454 cells. **A)** Osteocyte-like cells, OCY454, were differentiated for 12 days. Cells were combinatorially treated with inhibitor verteporfin (VP) and 5 ng/ml TGF-β1 at day 21. **B-G)** mRNA expression, normalized to 18s rRNA, was evaluated for **(B)** serpin family E member 1 (SerpinE1), **(C)** cysteine-rich angiogenic inducer 61 (Cyr61), **(D)** connective tissue growth factor (Ctgf), **(E)** cathepsin K (Ctsk, **(F)** matrix metalloproteinase-13 (Mmp13), and **(G)** matrix metalloproteinase-14 (Mmp14). Relative expression was expressed as fold vs. vehicle (PBS + DMSO)-treated cells. Sample sizes, N = 3.

## REFERENCES

1. Schuit SC., et al. (2004) Fracture incidence and association with bone mineral density in elderly men and women: the Rotterdam Study. Bone 34(1):195–202.

2. Delmas P, Li Z, Cooper C (2003) Relationship Between Changes in Bone Mineral Density and Fracture Risk Reduction With Antiresorptive Drugs: Some Issues With Meta-Analyses. J Bone Miner Res 19(2):330–337.

3. Sarkar S, et al. (2002) Relationships Between Bone Mineral Density and Incident Vertebral Fracture Risk with Raloxifene Therapy. J Bone Miner Res 17(1):1–10.

4. Hernandez CJ, Keaveny TM (2006) A biomechanical perspective on bone quality. Bone 39(6):1173–81.

5. Tranquilli Leali P, et al. (2009) Bone fragility: current reviews and clinical features. Clin Cases Miner Bone Metab 6(2):109–13.

6. Recker R, Lappe J, Davies KM, Heaney R (2004) Bone Remodeling Increases Substantially in the Years After Menopause and Remains Increased in Older Osteoporosis Patients. J Bone Miner Res 19(10):1628–1633.

7. Prideaux M, Findlay DM, Atkins GJ (2016) Osteocytes: The master cells in bone remodelling. Curr Opin Pharmacol 28:24–30.

8. Franz-Odendaal TA, Hall BK, Witten PE (2006) Buried alive: How osteoblasts become osteocytes. Dev Dyn 235(1):176–190.

9. Palumbo C, Palazzini S, Zaffe D, Marotti G (1990) Osteocyte Differentiation in the Tibia of Newborn Rabbit: An Ultrastructural Study of the Formation of Cytoplasmic Processes. Cells Tissues Organs 137(4):350–358.

10. Knothe Tate ML, Adamson JR, Tami AE, Bauer TW (2004) The osteocyte. Int J Biochem Cell Biol 36(1):1–8.

11. Buenzli PR, Sims NA (2015) Quantifying the osteocyte network in the human skeleton. Bone 75:144–150.

12. Hattner R, Epker BN, Frost HM (1965) Suggested sequential mode of control of changes in cell behaviour in adult bone remodelling. Nature 206(983):489–90.

13. Marotti G, Ferretti M, Muglia MA, Palumbo C, Palazzini S (1992) A quantitative evaluation of osteoblast-osteocyte relationships on growing endosteal surface of rabbit tibiae. Bone 13(5):363–8.

14. Baylink D, Wergedal J (1971) Bone formation by osteocytes. Am J Physiol Content 221(3):669–678.

15. Qing H, Bonewald LF (2009) Osteocyte Remodeling of the Perilacunar and Pericanalicular Matrix. Int J Oral Sci 1(2):59–65.

16. Qing H, et al. (2012) Demonstration of osteocytic perilacunar/canalicular remodeling in mice during lactation. J Bone Miner Res 27(5):1018–29.

17. Kawaki H, et al. (2011) Differential roles of CCN family proteins during osteoblast differentiation: Involvement of Smad and MAPK signaling pathways. Bone 49(5):975–989.

18. Zhao G, et al. (2018) CYR61/CCN1 Regulates Sclerostin Levels and Bone Maintenance. J Bone Miner Res. doi:10.1002/jbmr.3394.

19. Su J-L, et al. (2010) CYR61 regulates BMP-2-dependent osteoblast differentiation through the {alpha}v{beta}3 integrin/integrin-linked kinase/ERK pathway. J Biol Chem 285(41):31325–36.

20. Chen C-Y, et al. (2014) CCN1 induces oncostatin M production in osteoblasts via integrin-dependent signal pathways. PLoS One 9(9):e106632.

21. Si W, et al. (2006) CCN1/Cyr61 is regulated by the canonical Wnt signal and plays an important role in Wnt3A-induced osteoblast differentiation of mesenchymal stem cells. Mol Cell Biol 26(8):2955–64.

22. Smerdel-Ramoya A, Zanotti S, Deregowski V, Canalis E (2008) Connective tissue growth factor enhances osteoblastogenesis in vitro. J Biol Chem 283(33):22690–9.

23. Crockett JC, et al. (2007) The Matricellular Protein CYR61 Inhibits Osteoclastogenesis by a Mechanism Independent of α _v_ β _3_ and α _v_ β _5_. Endocrinology 148(12):5761–5768.

24. Zhao B, et al. (2008) TEAD mediates YAP-dependent gene induction and growth control. Genes Dev 22(14):1962–1971.

25. Zhang H, et al. (2009) TEAD transcription factors mediate the function of TAZ in cell growth and epithelial-mesenchymal transition. J Biol Chem 284(20):13355–62.

26. Yu F-X, Zhao B, Guan K-L (2015) Hippo Pathway in Organ Size Control, Tissue Homeostasis, and Cancer. Cell 163(4):811–828.

27. Mo J-S, Park HW, Guan K-L (2014) The Hippo signaling pathway in stem cell biology and cancer. EMBO Rep 15(6):642–56.

28. Zhang H, Pasolli HA, Fuchs E (2011) Yes-associated protein (YAP) transcriptional coactivator functions in balancing growth and differentiation in skin. Proc Natl Acad Sci U S A 108(6):2270–5.

29. Dole NS, et al. (2017) Osteocyte-Intrinsic TGF-β Signaling Regulates Bone Quality through Perilacunar/Canalicular Remodeling. Cell Rep 21(9):2585–2596.

30. Qin Z, Xia W, Fisher GJ, Voorhees JJ, Quan T (2018) YAP/TAZ regulates TGF-β/Smad3 signaling by induction of Smad7 via AP-1 in human skin dermal fibroblasts. Cell Commun Signal 16(1):18.

31. Miranda MZ, et al. (2017) TGF-β1 regulates the expression and transcriptional activity of TAZ protein via a Smad3-independent, myocardin-related transcription factor-mediated mechanism. J Biol Chem 292(36):14902–14920.

32. Varelas X, et al. (2008) TAZ controls Smad nucleocytoplasmic shuttling and regulates human embryonic stem-cell self-renewal. Nat Cell Biol 10(7):837–848.

33. Varelas X, et al. (2010) The Crumbs complex couples cell density sensing to Hippo-dependent control of the TGF-β-SMAD pathway. Dev Cell 19(6):831–44.

34. Delgado-Calle J, et al. (2017) Control of Bone Anabolism in Response to Mechanical Loading and PTH by Distinct Mechanisms Downstream of the PTH Receptor. J Bone Miner Res 32(3):522–535.

35. Bivi N, et al. (2012) Cell autonomous requirement of connexin 43 for osteocyte survival: Consequences for endocortical resorption and periosteal bone formation. J Bone Miner Res 27(2):374–389.

36. Kegelman CD, et al. (2018) Skeletal cell YAP and TAZ combinatorially promote bone development. FASEB J 32(5):2706–2721.

37. O’Brien CA, et al. (2008) Control of Bone Mass and Remodeling by PTH Receptor Signaling in Osteocytes. PLoS One 3(8):e2942.

38. Xiong J, et al. (2015) Osteocytes, not Osteoblasts or Lining Cells, are the Main Source of the RANKL Required for Osteoclast Formation in Remodeling Bone. PLoS One 10(9):e0138189.

39. Rhee Y, et al. (2011) PTH receptor signaling in osteocytes governs periosteal bone formation and intracortical remodeling. J Bone Miner Res 26(5):1035–1046.

40. Kegelman CD, et al. (2017) Skeletal cell YAP and TAZ redundantly promote bone development by regulation of collagen I expression and organization. bioRxiv:143982.

41. Tang SY, Herber R-P, Ho SP, Alliston T (2012) Matrix metalloproteinase-13 is required for osteocytic perilacunar remodeling and maintains bone fracture resistance. J Bone Miner Res 27(9):1936–50.

42. Jáuregui EJ, et al. (2016) Parallel mechanisms suppress cochlear bone remodeling to protect hearing. Bone 89:7–15.

43. Ploton D, et al. (1986) Improvement in the staining and in the visualization of the argyrophilic proteins of the nucleolar organizer region at the optical level. Histochem J 18(1):5–14.

44. Zambonin Zallone AZ, Teti A, Nico B, Primavera M V (1982) Osteoplastic activity of mature osteocytes evaluated by H-proline incorporation. Basic Appl Histochem 26(1):65–7.

45. Zambonin Zallone A, Teti A, Primavera M V, Pace G (1983) Mature osteocytes behaviour in a repletion period: the occurrence of osteoplastic activity. Basic Appl Histochem 27(3):191–204.

46. Holmbeck K, et al. (2005) The metalloproteinase MT1-MMP is required for normal development and maintenance of osteocyte processes in bone. J Cell Sci 118(1):147–156.

47. Liu-Chittenden Y, et al. (2012) Genetic and pharmacological disruption of the TEAD-YAP complex suppresses the oncogenic activity of YAP. Genes Dev 26(12):1300–5.

48. Kalajzic I, et al. (2004) Dentin matrix protein 1 expression during osteoblastic differentiation, generation of an osteocyte GFP-transgene. Bone 35(1):74–82.

49. Woo SM, Rosser J, Dusevich V, Kalajzic I, Bonewald LF (2011) Cell line IDG-SW3 replicates osteoblast-to-late-osteocyte differentiation in vitro and accelerates bone formation in vivo. J Bone Miner Res 26(11):2634–46.

50. Spatz JM, et al. (2015) The Wnt Inhibitor Sclerostin Is Up-regulated by Mechanical Unloading in Osteocytes in Vitro. J Biol Chem 290(27):16744–58.

51. Bartholin L, Wessner LL, Chirgwin JM, Guise TA (2007) The human Cyr61 gene is a transcriptional target of transforming growth factor beta in cancer cells. Cancer Lett 246(1–2):230–236.

52. Grotendorst GR (1997) Connective tissue growth factor: a mediator of TGF-beta action on fibroblasts. Cytokine Growth Factor Rev 8(3):171–9.

53. Pan J-X, et al. (2018) YAP promotes osteogenesis and suppresses adipogenic differentiation by regulating β-catenin signaling. Bone Res 6(1):18.

54. Xiong J, Almeida M, O’Brien CA (2018) The YAP/TAZ transcriptional co-activators have opposing effects at different stages of osteoblast differentiation. Bone 112:1–9.

55. Bellido T (2014) Osteocyte-driven bone remodeling. Calcif Tissue Int 94(1):25–34.

56. Nishida T, Emura K, Kubota S, Lyons KM, Takigawa M (2011) CCN family 2/connective tissue growth factor (CCN2/CTGF) promotes osteoclastogenesis via induction of and interaction with dendritic cell-specific transmembrane protein (DC-STAMP). J Bone Miner Res 26(2):351–363.

57. Canalis E, Zanotti S, Beamer WG, Economides AN, Smerdel-Ramoya A (2010) Connective Tissue Growth Factor Is Required for Skeletal Development and Postnatal Skeletal Homeostasis in Male Mice. Endocrinology 151(8):3490–3501.

58. Li CY, et al. (2006) Mice Lacking Cathepsin K Maintain Bone Remodeling but Develop Bone Fragility Despite High Bone Mass. J Bone Miner Res 21(6):865–875.

59. Hiemer SE, Szymaniak AD, Varelas X (2014) The transcriptional regulators TAZ and YAP direct transforming growth factor β-induced tumorigenic phenotypes in breast cancer cells. J Biol Chem 289(19):13461–74.

60. Wei H, et al. (2017) Hypoxia induces oncogene yes-associated protein 1 nuclear translocation to promote pancreatic ductal adenocarcinoma invasion via epithelial–mesenchymal transition. Tumor Biol 39(5):101042831769168.

61. Leivonen S-K, Chantry A, Häkkinen L, Han J, Kähäri V-M (2002) Smad3 Mediates Transforming Growth Factor-β-induced Collagenase-3 (Matrix Metalloproteinase-13) Expression in Human Gingival Fibroblasts. J Biol Chem 277(48):46338–46346.

62. Vashishth D, Verborgt O, Divine G, Schaffler MB, Fyhrie DP (2000) Decline in osteocyte lacunar density in human cortical bone is associated with accumulation of microcracks with age. Bone 26(4):375–380.

63. Bonewald LF (2006) Mechanosensation and Transduction in Osteocytes. Bonekey Osteovision 3(10):7–15.

64. Dupont S, et al. (2011) Role of YAP/TAZ in mechanotransduction. Nature 474(7350):179–83.

65. Gardinier JD, Al-Omaishi S, Morris MD, Kohn DH (2016) PTH signaling mediates perilacunar remodeling during exercise. Matrix Biol 52–54:162–175.

66. Dyment NA, et al. (2016) High-Throughput, Multi-Image Cryohistology of Mineralized Tissues. J Vis Exp (115):e54468–e54468.

67. Arganda-Carreras I, Fernández-González R, Muñoz-Barrutia A, Ortiz-De-Solorzano C (2010) 3D reconstruction of histological sections: Application to mammary gland tissue. Microsc Res Tech 73(11):1019–1029.

68. Mader KS, Schneider P, Müller R, Stampanoni M (2013) A quantitative framework for the 3D characterization of the osteocyte lacunar system. Bone 57(1):142–154.

69. Hemmatian H, et al. (2018) Age-related changes in female mouse cortical bone microporosity. Bone 113:1–8.

70. Tiede-Lewis LM, et al. (2017) Degeneration of the osteocyte network in the C57BL/6 mouse model of aging. Aging (Albany NY) 9(10):2190–2208.

71. Heveran CM, Rauff A, King KB, Carpenter RD, Ferguson VL (2018) A new open-source tool for measuring 3D osteocyte lacunar geometries from confocal laser scanning microscopy reveals age-related changes to lacunar size and shape in cortical mouse bone. Bone 110:115–127.

72. McCreadie BR, Hollister SJ, Schaffler MB, Goldstein SA (2004) Osteocyte lacuna size and shape in women with and without osteoporotic fracture. J Biomech 37(4):563–572.

73. Xin M, et al. (2013) Hippo pathway effector Yap promotes cardiac regeneration. Proc Natl Acad Sci U S A 110(34):13839–44.

74. Schneider P, Voide R, Stampanoni M, Donahue LR, Müller R (2013) The importance of the intracortical canal network for murine bone mechanics. Bone 53(1):120–128.

75. Akaike H (1974) A new look at the statistical model identification. IEEE Trans Automat Contr 19(6):716–723.

